# Aberrant Clonal Hematopoiesis Following Lentiviral Transduction of HSPC in a Rhesus Macaque

**DOI:** 10.1101/525691

**Authors:** Diego A Espinoza, Xing Fan, Di Yang, Stefan F Cordes, Lauren L Truitt, Katherine R Calvo, Idalia M Yabe, Selami Demirci, Kristin J Hope, So Gun Hong, Allen Krouse, Mark Metzger, Aylin Bonifacino, Rong Lu, Naoya Uchida, John F Tisdale, Xiaolin Wu, Suk See DeRavin, Harry L Malech, Robert E Donahue, Chuanfeng Wu, Cynthia E Dunbar

**Author notes:** Equal contributions (Diego A Espinoza, Xing Fan, Di Yang). Corresponding authors Chuanfeng Wu, Translational Stem Cell Biology Branch, NHLBI, NIH, Bethesda, Maryland 20892, Cynthia Dunbar, Translational Stem Cell Biology Branch, NHLBI, NIH, Bethesda, Maryland 20892.

## Abstract

Lentiviral vectors (LV) have been used for the delivery of genes into hematopoietic stem and progenitor cells (HSPC) in clinical trials worldwide. LV, in contrast to retroviral vectors, have not been associated with insertion site-associated malignant clonal expansions, and thus have been considered safer. Here, however, we present a case of markedly abnormal dysplastic clonal hematopoiesis impacting the erythroid, myeloid and megakaryocytic lineages in a rhesus macaque transplanted with HSPCs that were transduced with a LV containing a strong retroviral murine stem cell virus (MSCV) constitutive promoter-enhancer in the LTR. 9 insertions were mapped in the abnormal clone, resulting in overexpression and aberrant splicing of several genes of interest, including the cytokine stem cell factor and the transcription factor PLAG1. This case represents the first clear link between a lentiviral insertion-induced clonal expansion and a clinically abnormal transformed phenotype following transduction of normal primate or human HSPC, and are thus concerning, and suggest that strong constitutive promoters should not be included within LV vectors.

## Introduction

Lentiviral vectors(LV) have been successfully used for the delivery of genetic material into hematopoietic stem and progenitor cells(HSPCs) in clinical trials for inherited non-malignant diseases^1-8^. In contrast to γ-retroviral vectors, which have shown to induce insertion-associated leukemias both in clinical trials^9^ and in non-human primate models^10^, LV have not been associated with malignant clonal expansions in clinical trials or large animal models^11^. LVs integrate preferentially within active genes rather than near transcriptional start sites^12,13^, and have thus been considered less likely to activate nearby proto-oncogenes. However, reports from recent clinical trials suggest that LVs are capable of inducing insertion-site associated clonal (albeit, non-malignant) expansions^8,14^. Long-term monitoring of transduced HSPC clones in human and non-human primates remains critical for assessing the safety of LV gene therapies.

Our laboratory has used a rhesus macaque (RM) genetic barcoding model to study the output of lentivirally-transduced transplanted HSPC at a clonal level over time ^15-18^; such a setting closely resembles that of lentiviral HSPC gene therapy trials in which autologous HSPC are collected, transduced, and reinfused into the patient. In our published and unpublished studies, we have tracked over 101,000 individual lentiviral vector insertions in 9 transplanted macaques followed for up to six years and observed no clonal expansions associated with a transformed or aberrant hematologic phenotype. We now report on the development of aberrant, markedly dysplastic fatal clonal hematopoiesis in one animal, impacting the erythroid, myeloid and megakaryocytic lineages and originating from a single transduced HSPC clone containing 9 integrated LVs.

## Methods (See Supplemental Methods for additional details)

### Autologous transplantation of lentivirally-barcoded HSPC

CD34^+^ HSPC were collected from ZL34, a 4-year old male rhesus macaque, following mobilization with G-CSF/AMD1300, transduced(MOI=25) with a barcoded lentivirus library, and reinfused into the macaque following TBI as described^15-18^. ZL34 received weekly dosing with 5mg/kg cidofovir to suppress CMV reactivation.

### Barcode retrieval

DNA was extracted using the DNeasy Blood & Tissue Kit(Qiagen). 200ng was used for barcode retrieval PCR with Phusion High-Fidelity DNA Polymerase(ThermoFisher) via 28 cycles of 98°C-10 seconds, 70°C-30 seconds, 72°C-30 seconds; then 72°C-10 minutes. Indexed or non-indexed forward primers and indexed reverse primers were added(Table S6) for multiplex sequencing. Following gel purification, 15-24 multiplexed samples were used to create a DNA library for sequencing on an Illumina HiSeq2500/3000. Custom Python code(https://github.com/d93espinoza/barcode_extracter) was used for extraction and processing of barcodes from FASTQ files as described^15-18^. For modified barcode recovery, “Starcode” software(https://github.com/gui11aume/starcode) was used for extraction and processing of barcodes. Visualization was performed using custom R code. For original data, contact.

## Results

### Lentiviral transduction and transplantation

We developed a lentiviral barcoding model to quantitatively study the output of thousands of individual HSPC following autologous transplantation^15-18^. As part of our ongoing studies, we transplanted animal ZL34 with autologous CD34^+^ HSPC transduced with a lentiviral barcoding vector driving GFP expression via the strong murine retroviral MSCV promoter-enhancer inserted within the lentiviral long terminal repeat(LTR)(Figure 1A). The transduction/transplantation protocols utilized were unchanged from our prior experience^15-18^ and the multiplicity of infection(MOI), CD34^+^ transplanted dose(cells/kg), and GFP% of the infused cells were within the ranges utilized for prior animals(Table S1, n = 10). ZL34, along with another monkey, ZL40(Table S1), received CMV-suppressing cidofovir for 4 months to study the impact of CMV reactivation on immune reconstitution. With the exception of ZL34, all other animals, including ZL40 also receiving cidofovir on the same schedule, continue to show stable, highly polyclonal output from transduced HSPC with no evidence for genotoxicity or malignant clonal expansions at latest follow up(4 months - 70 months, median: 25 months), regardless of having received HSPC transduced with vectors containing either an internal EF1α promoter(n=3) or the LTR-embedded MSCV promoter(n= 7), as previously published^15-17^.

**Figure 1.**
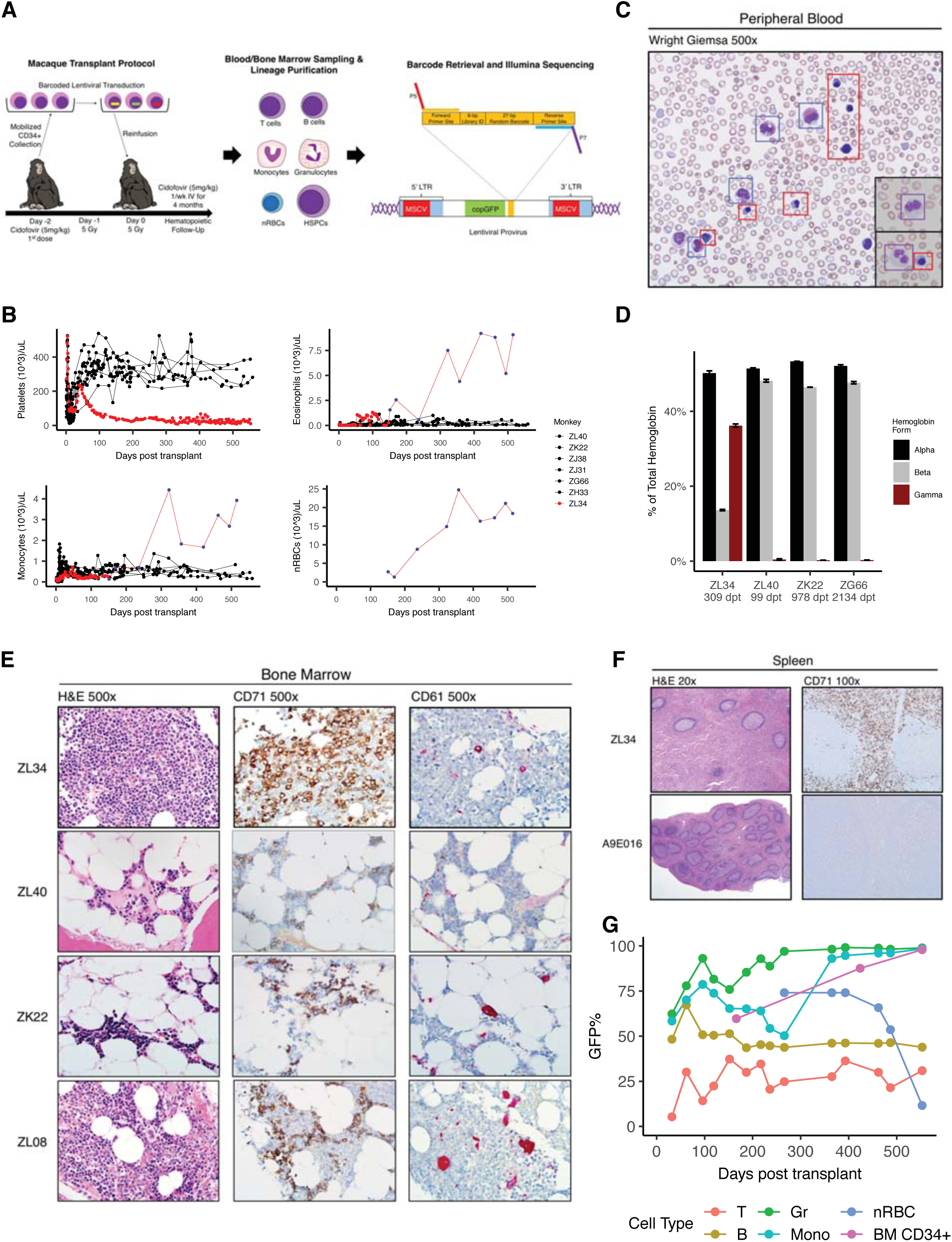
Rhesus macaque autologous transplantation model and development of aberrant hematopoiesis in rhesus macaque ZL34. **(A)** Transplantation timeline for ZL34. HSPC were mobilized with G-SCF and AMD3100, collected via apheresis, enriched for CD34^+^ HSPC, and transduced with a barcoded lentiviral vector. Following total body irradiation, transduced autologous CD34^+^ HSPC were reinfused, and cidofovir was administered. The integrated form of the provirus is shown, detailing position of the MSCV promoter/enhancer(s), copGFP marker gene, and high-diversity barcode library consisting of a 6bp library ID and a 27bp random nucleotides, flanked by sequencing primer binding sites. (**B)** Platelet, eosinophil, monocyte, and nRBC concentrations in the blood over time in seven macaques receiving cells transduced with the barcoded LV vector containing the MSCV promoter/enhancer, including ZL34. Counts were determined by Coulter counter(red or black circles) or by automated PB smear scanning with Cellavision DM(blue dots – ZL34). **(C)** Wright’s stained ZL34 PB smear from day 424 post-transplantation showing frequent eosinophils(blue squares), nucleated erythroid cells(red squares) which are not found in normal rhesus macaque blood, and hypogranular neutrophils with dysplastic nuclei(black insets, purple squares). **(D)** HPLC assay of hemoglobin chain composition in the blood partitioned into α, β,and γ globin molecules for ZL34 309 days post-transplantation(dpt) and three barcoded macaques without clonal expansions(ZL40, ZK22 and ZG66 99-2134 days post-transplantation). ZL34 values correspond to a predicted fetal hemoglobin of 70%. Error bars denote standard deviation for three technical replicates. **(E)** BM biopsies from ZL34(day 424), two healthy transplanted macaques(ZK22 and ZL40), and one healthy non-transplanted macaque (ZL08). H&E: hematoxylin and eosin; CD71: immunohistochemistry for the erythroid precursor-specific transferrin receptor (CD71); CD61: immunohistochemistry for the megakaryocyte-specific glycoprotein IIIa (CD61), note that ZL34’s megakaryocytes are small and uninuclear; **(F)** Spleen from ZL34 and a control transplanted macaque (A9E016). H&E: hematoxylin and eosin; CD71: immunohistochemistry for the erythroid precursor-specific transferrin receptor(CD71). **(G) %**GFP^+^ cells by flow cytometry over time in ZL34 PB T cells, B cells, granulocytes, monocytes, and nRBC, and BM CD34^+^ cells.

### Development of abnormal hematopoiesis

ZL34 engrafted normally post-transplantation as compared to other animals, promptly recovering neutrophils to greater than 500/uL, platelets to >200,000 platelets/uL, and hemoglobin to >9 g/dL(Figures 1B; S1A, B). However, ZL34’s platelet count declined beginning 49 days post-transplantation to levels far below the range maintained in other transplanted macaques(Figure 1B), requiring platelet transfusions to treat bleeding. Waxing and waning eosinophilia, without identified infectious, drug or allergic etiology, emerged 50 days post-transplantation and became persistent by day 300(Figure 1B). Eosinophilia has not developed in other macaques on the same transplant regimen. The morphology of neutrophils became markedly dysplastic, with hypogranulartiy and bilobulated/hypolobulated nuclei, despite normal neutrophil counts and lack of circulating blasts(Figures 1C; S1A). Monocytes and basophils also rose to well above normal range by day 300(Figures 1B; S1C). In contrast, lymphocyte counts remained within expected post-transplant ranges(Figure S1D).

Nucleated red blood cells (nRBCs) are normally found only in the bone marrow(BM) of non-human primates and humans, with nuclear extrusion occurring prior to release into the blood. Strikingly, large numbers of nRBCs were prematurely released into the blood in ZL34, first noted on day 151 and reaching concentrations greater than 25,000/µL(Figures 1B-C). No other transplanted macaques have had detectable circulating nRBCs. ZL34’s hemoglobin levels and red blood cell counts remained within the normal range until one year post-transplantation(Figure S1B,E). Analysis of globin chain ratios showed marked upregulation of γ-globin chains(and thus, fetal hemoglobin) in ZL34’s red blood cells long-term, in contrast to the normal dominance of adult β chains by day 100 or later in other transplanted macaques^19^(Figure 1D).

Compared to healthy transplanted(ZL40 and ZK22) and non-transplanted(ZL08) RMs, ZL34’s BM at day 424 was hypercellular with increased eosinophils and a marked erythroid dominance, and markedly dysplastic uninuclear micro-megakaryocytes(Figure 1E). Blasts were not increased. Day 543 BM karyotype was normal in 20/20 metaphases. Splenic red pulp was markedly expanded due to extramedullary hematopoiesis, with infiltration of immature CD71^+^ erythroid cells(Figure 1F), an abnormal finding in macaques or humans. Pathologically and clinically, this hematologic disorder would be classified by WHO criteria as an overlap myelodysplastic/ myeloproliferative neoplasm with extramedullary hematopoiesis. Bleeding necessitated euthanasia of ZL34 on day 553.

Inclusion of copGFP in the vector allowed analysis of the level of engraftment with transduced cells. The %GFP^+^ T and B cells remained generally stable over time and within the upper range of values observed in other barcoded RMs^15-18^ (Figure 1G, S1F). In contrast, %GFP^+^ granulocytes(including neutrophils and eosinophils) and monocytes approached 100%, coincident with development of markedly abnormal blood counts, suggesting expansion of transduced cells within these lineages. %GFP expression in the circulating nRBCs was also very high, but dropped precipitously before euthanasia, potentially due to transgene silencing. Other barcoded animals did not show expansion of GFP+ cells of any lineage over time, with similar levels in all lineages other than delayed reconstitution of GFP+ T cells due to TBI effects on the thymus(Figure S1F and^15-17^).

The %GFP^+^ of CD34^+^ BM HSPC also increased over time, reaching over 90%. Flow cytometric analysis of CD34^+^ subsets based on prior validation in macaques^20^, showed that the most primitive CD34^**+**^CD90^**+**^CD45RA^**-**^ population, highly-enriched for multipotent long-lived HSCs, was most markedly GFP^**+**^, with lower percentages in the CD34^**+**^CD90^−^CD45RA^**+**^ population, containing lymphoid precursors^20^(Figure S2A-C). Of note, the CD34^**+**^CD90^**+**^CD45RA^**-**^ population was also markedly-expanded in ZL34 BM compared to normal, and present in the spleen, suggesting a differentiation block as well as extramedullary hematopoiesis(Figure S2A, B). These findings are consistent with development of an MPN/MDS neoplastic syndrome resulting from abnormalities of transduced primitive HSPC.

### A single expanded transduced clone was responsible for abnormal hematopoiesis

We performed quantitative barcode retrieval from T cells, B cells, neutrophils, eosinophils, monocytes, nRBCs and CD34+ cells using primers flanking the barcode region^15-18^(Figure 2A-G). 6 barcodes contributed equally and accounted for 100% of barcodes from nRBCs at all time points. The same 6 clones appeared and expanded in neutrophils and monocytes beginning at day 187, replacing earlier highly polyclonal contributions. Eosinophils at day 494 contained only the same 6 dominant barcodes. In contrast, T and B cells maintained polyclonal diversity over time, with intermittent detection of much lower levels of the 6 barcodes, likely due to contamination with clonally-expanded myeloid or erythroid cells during sorting, and/or aberrant marker expression on dysplastic myeloid cells.

**Figure 2.**
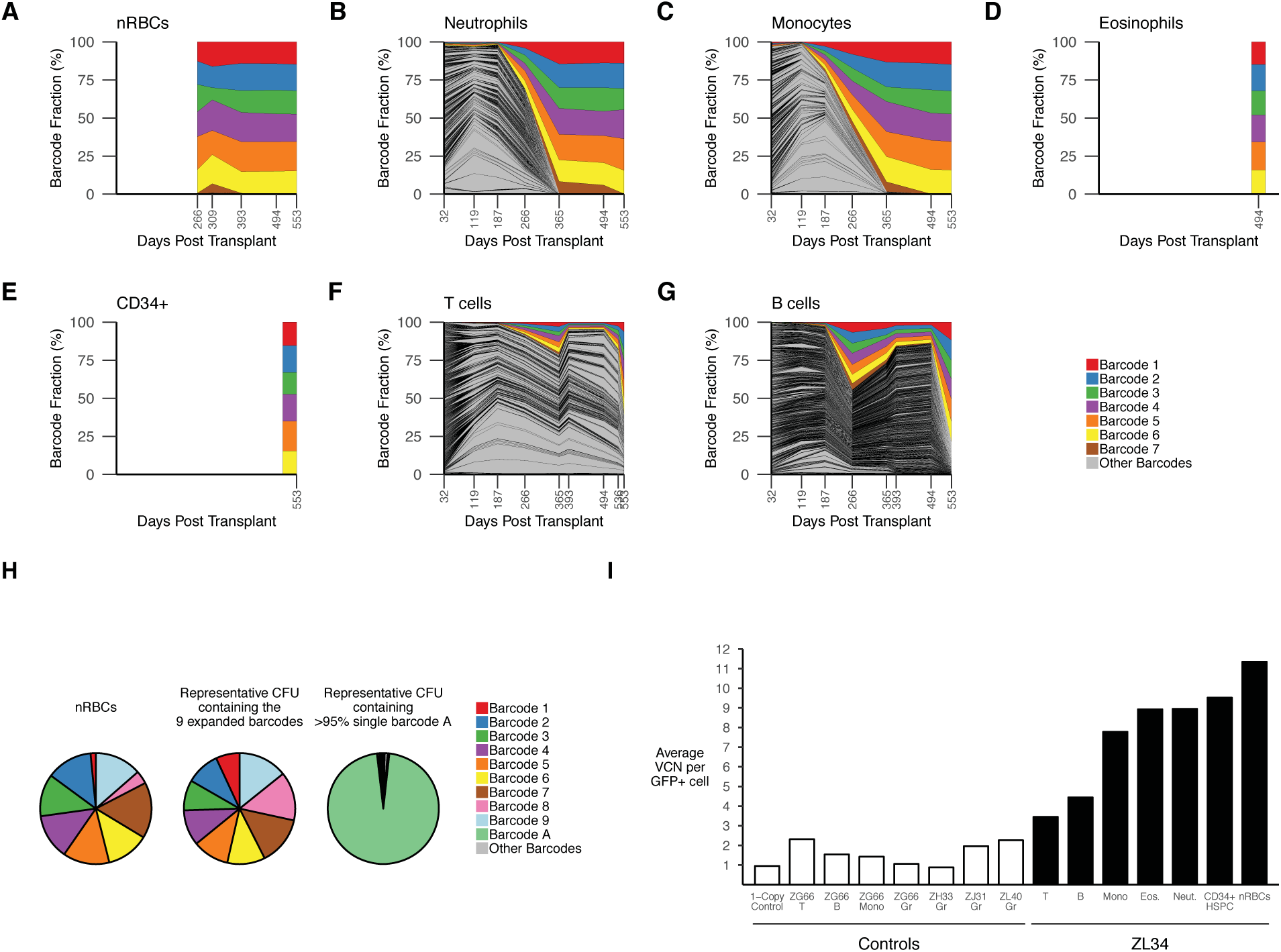
Clonal tracking via barcode retrieval in ZL34. Longitudinal clonal tracking of barcodes retrieved from nRBC**(A)**, neutrophils**(B)**, monocytes**(C)**, eosinophils**(D)**, BM CD34+ cells**(E)**, T cells**(F)**, and B cells**(G)**. Each graph shows the shows fractional contributions from individual barcodes over time, with each colored ribbon representing the expanded barcodes detected via standard barcode recovery and all other barcodes shaded in gray. **(H)**Modified barcode recovery using the alternative forward primer on ZL34 nRBCs(393 days post-transplant) showing retrieval 9 expanded barcodes. Modified barcode recovery using the same primers was performed on single-cell derived GFP+ CFU from ZL34 BM CD34^+^ cells(266 days post-transplant) plated at low density. Representative barcode analysis on a myeloid CFU containing all 9 barcodes is shown, matching those found in the circulating nRBC, confirming integration of the 9 barcodes in one original HSPC. For comparison, barcode analysis on a myeloid CFU from the same plate found to contain a single barcode, Barcode A, is shown, confirming lack of cellular contamination across the plate that would complicate analysis. The small sector shown in grey represents multiple other barcodes with very low read counts likely resulting from residual single cells not forming CFU. **(I)** Vector copy number per cell quantified via quantitative PCR on Gr(granulocytes), Mono(monocytes), T cells, B cells, Eos(eosinophils), nRBC and CD34^+^ BM HSPC from ZL34(black bars, days 494-553 post-transplant) and additional barcoded transplanted macaques(white bars, days 278-1085 post-transplant).

A 7^th^ barcode was intermittently detected within the abnormal lineages, due to a single mutation-related mismatch at the 3’ end of the primer annealing site (Figure 2A-G), prompting redesign of an upstream primer to retrieve any additional insertions. Using this strategy, we recovered an additional 2 barcodes from circulating nRBCs, for a total of 9(Figure S3).

Because these expanded barcodes contributed roughly equally to every sample, we suspected multiple vector insertions within a single originating clonal HPSC. We plated ZL34 and control BM CD34^+^ cells in semi-solid media allowing barcode retrieval from individual colony-forming units (CFU). Of note, despite the erythroid predominance in the BM, a reduced ratio of erythroid/myeloid colonies was observed in ZL34 compared to normal BM(Figure S4). We plucked individual GFP+ myeloid CFU. All 9 barcodes were retrieved together from the majority of individual CFU(Figure 2H). Rare CFU lacked the 9 barcodes of interest but each instead contained 1-4 other barcodes.

Vector copy number (VCN) in samples from ZL34 and other lentivirally-barcoded RMs was determined by quantitative PCR. The normalized VCN per GFP^+^ cell for nRBCs, eosinophils, monocytes, neutrophils and CD34^+^ HSPC from late time points in ZL34 averaged 8.5 copies/cell (Figure 2I) which along with the CFU barcode analysis supports a conclusion that expansion from a single HSPC clone with 9 vector insertions was responsible for the myelodysplastic/myeloproliferative neoplasm in ZL34. The uninvolved B and T cells lineages in ZL34 had much lower VCN per GFP^+^ cell, as did myeloid and lymphoid lineages in other barcoded macaques(Figure 2I).

### Retrieval of LV insertion sites for the 9 dominant barcodes

We recovered 9 LV insertions from circulating nRBCs using standard methodologies(Figure 3A;Table S2). In contrast, insertions retrieved from neutrophils at day 96 were polyclonal(Table S2). We were able to link each of the 9 dominant barcodes to one of the 9 nRBC insertions via PCR amplification and sequencing(Figure 3A). 4 insertions localized to introns within the *NCAM2, EIF3E, PLAG1, and IMMP2L* genes, with *PLAG1, NCAM2* and *EIF3E* proviral insertions in the same orientation as endogenous transcripts, and the *IMMP2L* insertion in the opposite orientation. A number of additional genes were located within 1 Mb windows surrounding each of the 9 insertions(Figures 3B;S5). One insertion was less than 20 kb downstream of *KITLG,* encoding stem cell factor(*SCF*), a cytokine of importance in hematopoiesis. None of the insertions localized to known CTCF binding sites, or rhesus macaque sequences homologous to open chromatin regions mapped in human hematopoietic cells(Figure S6).

**Figure 3.**
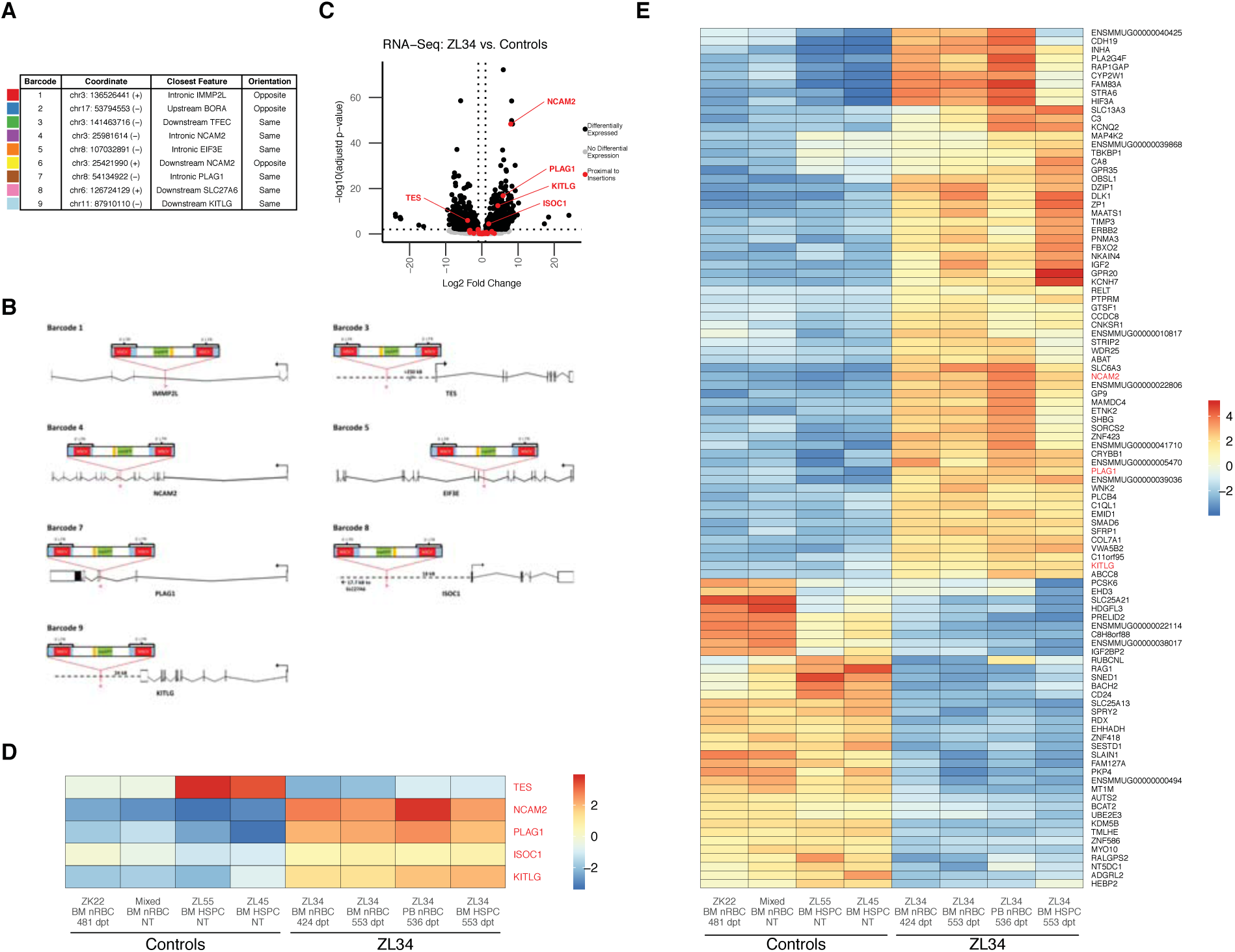
Identification of insertion sites and impact on gene expression in hematopoietic cells. **(A)** Localization in the rhesus genome, closest gene, and proviral orientation relative to the closest gene for the 9 insertion sites corresponding to the 9 expanded barcodes. **(B)** Orientation and location of the provirus relative to the 5 proximally differentially expressed genes and 2 genes with intronic disruptions. **(C)** Volcano plot of RNA-seq data from normal control vs ZL34 nRBC and CD34+ cells showing all differentially expressed genes (black) as defined by padj < 0.01 and log2(fold change) > 1 or log2(fold change) < −1. Non-differentially expressed genes shown in grey. Differentially-expressed genes within 1 Mb upstream to 1 Mb downstream of any of the 9 insertion sites are colored in red and labelled. Two control BM nRBC samples (one from transplanted macaque ZK22, one pooled from two non-transplanted macaques) and two control BM CD34+ samples (from two different non-transplanted macaques) were analyzed along with two independent BM nRBC samples, one PB nRBC sample, and one CD34+ sample from ZL34**. (D)** Mean-centered gene expression levels of *NCAM2, KITLG, PLAG1, ISOC1*, and *TES*. Expression levels are rlog values from DESeq2. NT = non-transplanted, dpt = days post-transplant. **(E)** Mean-centered gene expression levels of the top 100 differentially-expressed genes (by adj. p-value) in ZL34 compared to control nRBC and CD34^+^ HSPC samples. Expression levels are rlog values from DESeq2. A complete list of differentially-expressed genes is given in Table S3.

### Impact of insertions on gene expression

We performed bulk RNA-seq on ZL34 PB nRBCs, BM nRBCs, and BM CD34^+^ HSPCs obtained on days 424-553 post-transplant, as well as on control RM BM nRBCs and BM CD34^+^ HSPCs. No control RM PB nRBCs were obtained, since nRBCs are not found in normal PB. We asked whether expression of genes interrupted by or within 1Mb of any of the 9 insertions(Figures S5,S7) were differentially-expressed in ZL34 compared to controls. We found 5 genes within these windows to be significantly differentially expressed combining nRBC and CD34 data, regressing out the source(PB or BM) and cell type(HSPC or nRBC) such that only significant differences attributed to condition(ZL34 cells vs controls) were identified(log2 fold change > 1 or < −1, and adjusted p-value < 0.01)(Figure 3C-D). *NCAM2* and *PLAG1* with intronic insertions in the same orientation were overexpressed, *KITLG* with an insertion 24kB downstream was overexpressed*, ISOC1* with an insertion 18kB upstream was overexpressed, and *TES* with an insertion 250kB upstream was under-expressed. When restricting analysis to nRBCs, 4 genes were differentially expressed: *NCAM2*, *PLAG1, KITLG, and ISOC1*(Figure S8A-B).

When considering all recovered transcripts whether adjacent to insertions or not, a number of additional genes were differentially expressed. *KITLG, NCAM2, and PLAG1* were among the top 100 in combined HSPC and nRBC analysis(Figure 3E; Table S3) and were either within the top 100 or differentially expressed within nRBCs alone(Figure S8C;Table S3). As expected from the hemoglobin analysis(Figure 1D), *HBG2* transcripts, encoding the γ-globin chain, were significantly upregulated in ZL34 nRBCs(Figure S8C).

SCF binds to the c-kit tyrosine kinase receptor on HSPC and stimulates survival and proliferation^21,22^. SCF is normally produced by BM stromal elements, not HSPC. We measured soluble SCF levels in ZL34 and control blood and in media conditioned by cultured ZL34 and normal PB MNC(Figure 4A). The secreted isoform of SCF was not detected above background in cultured samples. An increase in SCF was detected in ZL34 serum. Using an antibody that reacts with the transmembrane isoform of SCF, we found clear increased expression of cell-associated SCF in hematopoietic cells as shown by immunohistochemistry on marrow sections, and via FACS(Figures 4B-C; S9B).

**Figure 4.**
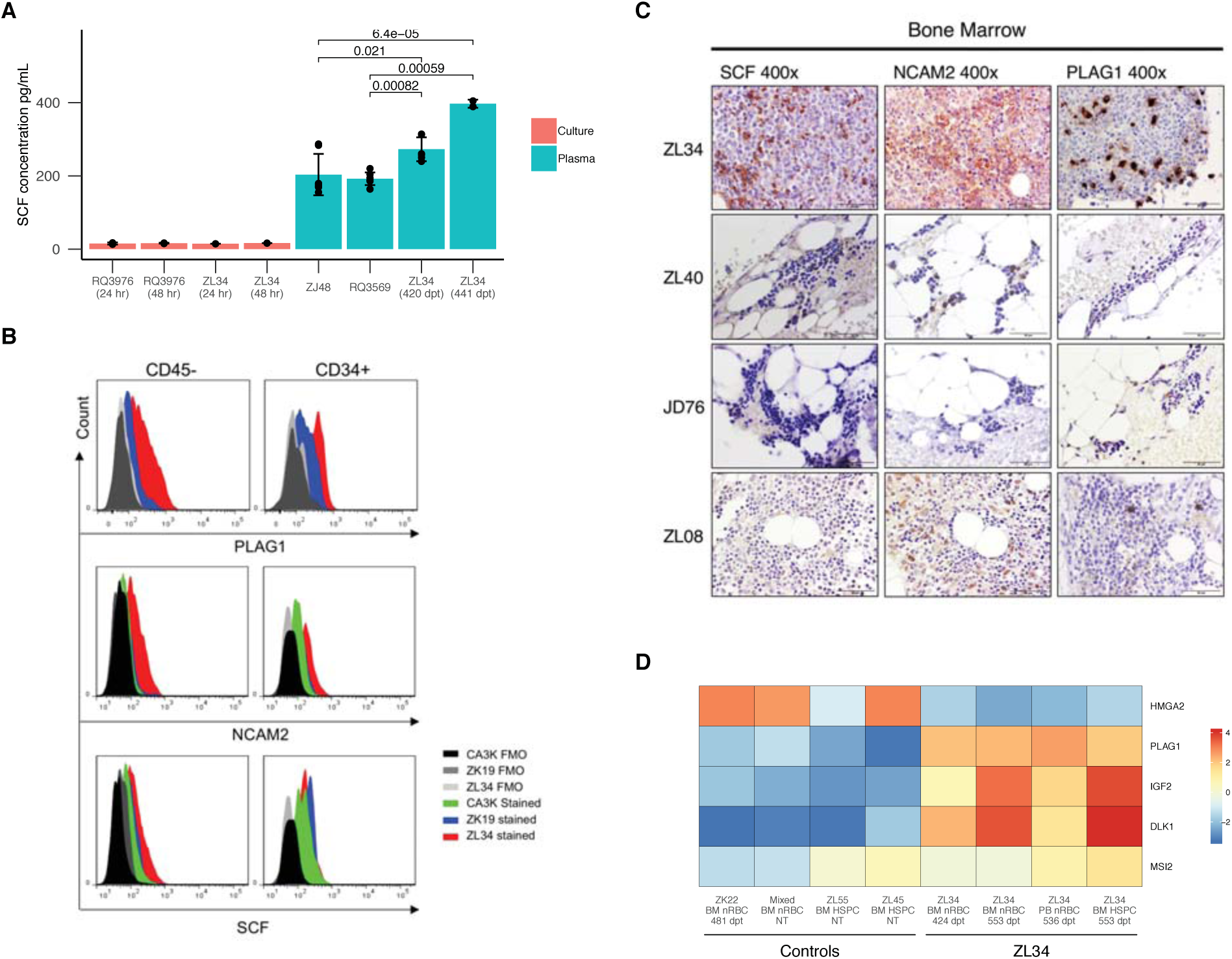
Analysis of dysregulated expression of *KITLG, NCAM2,* and *PLAG1*. **(A)** Stem cell factor (SCF) concentrations in serum and in culture media from high density cultured PB MNC, for ZL34 and controls as determined by ELISA. Error bars are shown for technical replicates. p-values are shown and determined by Welch’s two-sample t-test. **(B)** Expression of PLAG1, NCAM2, and SCF assessed by flow cytometry in ZL34 and control non-transplanted normal macaque (ZK19), and a macaque recovering from malarial anemia (CA3K). Gating of BM CD45^**-**^ cells were used as an alternative marker for nRBCs in these studies, due to destruction of the CD71 epitope during permeabilization for intracellular staining. >85% of CD45-cells are nRBCs (Fig.S6C). ZK19 is a non-transplanted normal monkey. **(C)** ZL34 and control BM samples (ZL08: non-transplanted, JD46 and ZL40: barcoded animals) stained via immunohistochemistry for PLAG1, SCF, and NCAM2. **(D)** Mean-centered gene expression levels for *PLAG1*, its putative upstream regulator *HMGA2*, and PLAG1 downstream targets *IGF2, DLK1*, and *MSI2* in ZL34 and control samples. Expression levels are rlog values from DESeq2.

*PLAG1* is a zinc finger transcription factor and proto-oncogene upregulated in salivary gland tumors^23^ and in murine and human myeloid leukemias^24,25^. At the protein level, PLAG1 was overexpressed on a per cell basis in ZL34 CD45^**-**^ nRBCs, HSPCs, and monocytes but not T or B cells, as shown via intracellular FACS and most strikingly immunohistochemistry, revealing numerous intensely PLAG1-positive BM cells compared to controls(Figure 4B,C; S9B). Increased activity of PLAG1 was confirmed by significant increases in expression of known downstream targets, including *IGF2*, which encodes insulin-like growth factor 2(p=3.66 * 10^−17^) and *DLK1* (p=7.70 * 10^−31^), both imprinted genes reported to be dysregulated in PLAG1 associated tumors^26-29^(Figure 4D). *MSI2*, a gene under direct regulation by PLAG1^30^ and whose product Musashi-2 is reported to promote HSPC expansion^31^ and self-renewal was also upregulated(p=5.37 * 10^−5^)(Figure 4D). Of note, mRNA encoding *HMGA2*, a transcription factor functionally upstream of PLAG1^32^, was expressed at a lower level in ZL34 nRBCs and HSPCs compared to control samples. Insertional overexpression of *HMGA2* has been previously-linked to clonal expansion in a lentiviral gene therapy clinical trial^8^, suggesting HMGA2 may stimulate HSPC clonal expansion via its impact on PLAG1 expression and activation of downstream targets active on HSPC such as *MSI2,* and that tonic low level HMGA2 expression may be involved in normal HSPC homeostasis, controlled by feedback from downstream targets.

*NCAM2* is an adhesion molecule required for cell-cell or cell-matrix interactions during nervous system development, but is not normally expressed in hematopoietic cells, and dysregulation has not been linked to hematologic disease. Both FACS and immunohistochemistry confirmed overexpression of NCAM2 at a protein level in CD34^+^ cells, nRBC, and monocytes(Figure 4B,C; S9B). Decreased expression of the tumor suppressor *TES* has not been previously linked to hematologic malignancies. The *TES* gene itself was not directly disrupted by an insertion, thus downregulation of this gene must have been indirect, via disruption of a regulatory element. There is little information about the gene product of *ISOC1*. It is not expressed in hematopoietic cells and has no known link to hematologic disease.

### Insertion-related aberrant splicing

LV vectors have been reported to disrupt splicing via formation of hybrid vector-gene transcripts expressed from viral promoters, resulting in dysregulated expression, alterations in isoform, and/or loss of transcript regulatory sequences^33,34^. We first interrogated RNA-seq data for any differentially-spliced genes and found that ZL34 *PLAG1* transcripts contained significantly lower levels of exons E1 and E2 compared to controls. We did not detect significant variable splicing for any other gene with intronic insertions(Table S4). However, several other genes of hematopoietic interest, including HIF3A, demonstrated both changes in isoform abundance(Table S4) and overall upregulation(Figure 3E). We next mined RNA-seq data for fusion transcripts. We detected fusions between vector and *PLAG1* sequences in ZL34 nRBC and abundant hybrid forms containing vector sequences spliced to E3, before the translation start site, including isoforms with and without E4(Figure 5B;Table S5), predicted to encode the two major isoforms of PLAG1 protein^35^. We also detected fusion *EIF3E* and *NCAM2* transcripts(Figure S10;Table S5), however, *EIF3E* was not overall upregulated in ZL34(Figure S7). A subset of these hybrid transcripts were confirmed by RT-PCR(Figures 5B;S10). We did not detect hybrid transcripts for the other genes with intronic insertions nor any non-LV fusion transcripts.

**Figure 5.**
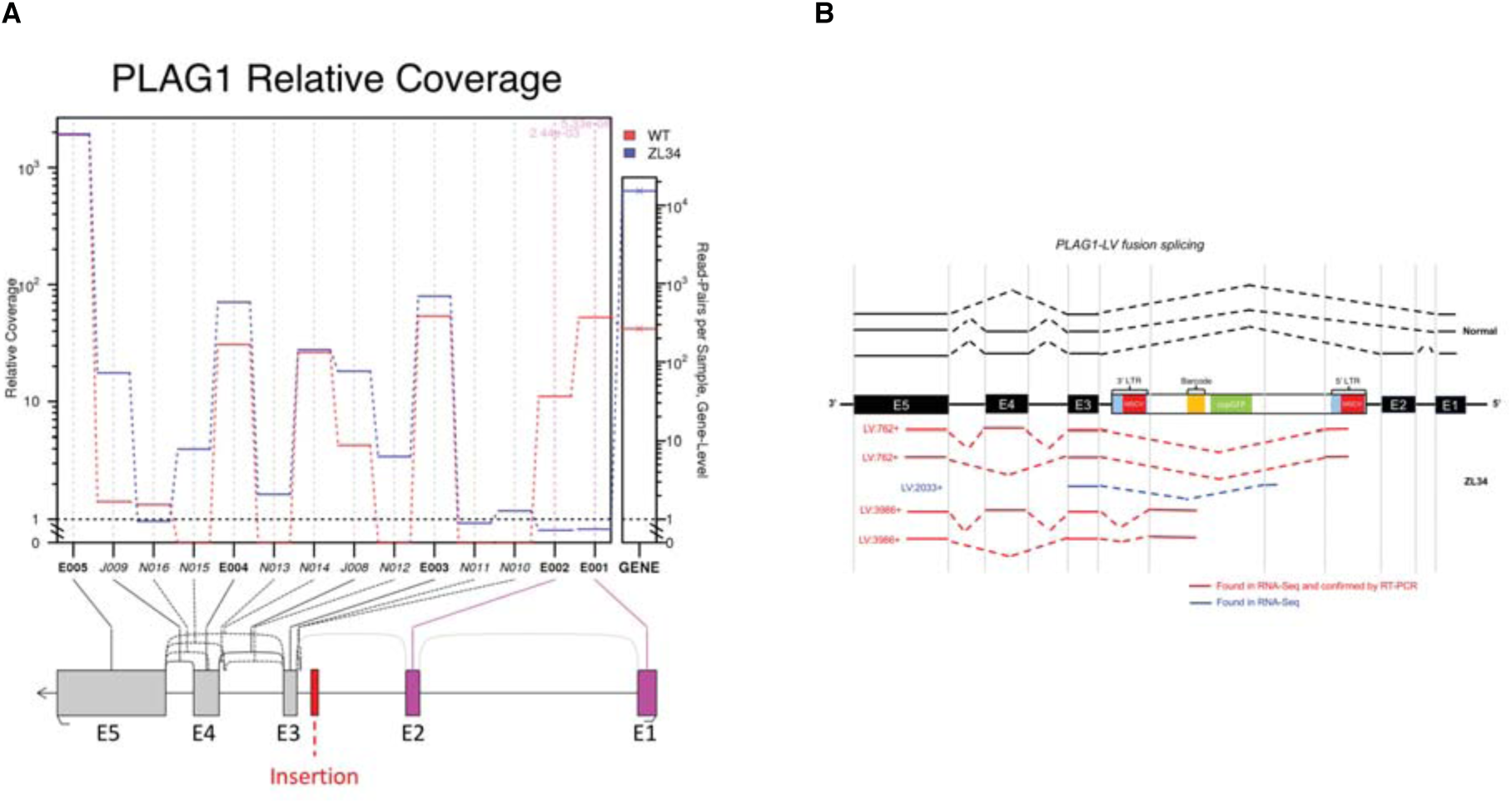
Abnormal splicing related to lentiviral insertions. **(A)** Differential expression of exons within ZL34 relative to control samples as determined from RNA-seq data, showing significant downregulation of exons 1 and 2 within ZL34 *PLAG1* transcripts relative to control macaque nRBCs and HSPCs. **(B)** Hybrid vector-*PLAG1* mRNA species detected via RNA-seq fusion transcript analysis and/or confirmation by PCR and Sanger sequencing in ZL34 and control nRBC. Lines denote transcripts detected in ZL34 (red) and normal controls (black). Results show absence of exons 1 and 2 in ZL34 PLAG1 transcripts, along with hybrid vector-*PLAG1* transcripts.

## Discussion

We describe the first case of lentiviral insertion-induced genotoxicity resulting in hematopoietic clonal expansion and a neoplastic phenotype in a large animal or human. Following autologous transplantation of HSPC transduced with a barcoded LV containing the MSCV promoter-enhancer within the LTR, one macaque developed a fatal myeloproliferative/myelodysplastic syndrome linked to clonal expansion from a single transduced HSPC.

The development of leukemias in clinical gene therapy trials and in primate models utilizing murine γ-retrovirus vectors(RV) to transduce HSPC stimulated a search for safer gene transfer systems^9-11^. Evidence implicated RV activating nearby cellular proto-oncogenes with strong viral enhancers, exacerbated by RV integration favoring transcription start sites(TSS)^13^. LV had endogenous enhancers removed, and were shown to have an integration pattern favoring gene bodies rather than promoters, hypothesized to be less likely to activate proto-oncogenes^36,37^. Genotoxicity models including *in vitro* immortalization of murine HSPC and acceleration of leukemia in tumor-prone mice predicted significantly lower genotoxicity from LV versus RV, even when strong viral promoters derived from RV were inserted within LV backbones^38-41^.

The majority of HSPC clinical gene therapy trials over the past decade have utilized LV to try and decrease the risk of genotoxicity and take advantage of increased efficiency of HSPC transduction associated with this vector class^1,42^. Most trials have utilized or are utilizing LV with internal weak constitutive or lineage-specific promoters, although several trials for diseases requiring very high transgene expression, such as adrenoleukodystrophy, continue to employ LV with very strong internal viral promoter/enhancers.

In the majority of our barcoded macaques, including ZL34, we used a LV construct containing the MSCV viral promoter/enhancer inserted within the LTR. The original MSCV RV vector was derived from a mutant murine myeloproliferative sarcoma virus (MSPV) with high expression in hematopoietic cells, and the MSCV promoter/enhancer was later shown to drive consistent high stable transgene expression in human HSPC when included in LV vectors^43,44^. In a previous study, we reported development of AML following transduction of macaque HSPC using an RV vector containing the MSCV promoter/enhancer^10^. However, in 6 macaques transplanted with HSPC transduced with the same LV barcoding vector containing the MSCV sequences, we have tracked over 65,000 individual clones for up to 5 years without clonal expansions. In addition, 3 macaques received HSPC transduced with a barcoded LV containing an internal EF1α promoter showed clonal patterns indistinguishable from the MSCV animals(Table S1 and^15^).

Because RV promoter/enhancers included in LV vectors promote high constitutive transgene expression, two clinical trials for adrenoleukodystrophy have included a RV promoter/enhancer similar to MSCV, termed MND, in the LV to drive high expression of the transgene^4,5,45,46^. In contrast to the vector utilized in ZL34, the adrenoleukodystrophy trial LV placed the RV promoter/enhancer internally, instead of within the LV LTR^4,5,45,46^. Tumor-prone mouse models have suggested that an internal strong promoter/enhancer is less genotoxic than the same promoter/enhancer placed within the LV LTR^39^. Thus far, no clonal expansions or hematopoietic toxicity have been reported in the 23 patients enrolled in these trials, with many years of follow-up. Other clinical trials using LV containing endogenous or weaker constitutive promoters have reported polyclonal hematopoiesis for years post-transplantation^6,47-50^. However, a thalassemia patient developed a marked clonal expansion linked to insertion of an LV containing an internal erythroid-specific promoter, resulting in increased expression and aberrant splicing of *HMGA2*, without any associated abnormalities of hematopoiesis^8^. Recently, persistence and massive clonal expansion of a CAR-T cell clone in a patient was linked to an LV insertion inactivating the epigenetic regulator TET2^14^.

The expanded neoplastic clone in our macaque had 9 insertions, with 4 out of the 9 insertions located within introns. We believe the most compelling candidate insertion driving the HSPC phenotype was within the gene encoding the zinc finger protein PLAG1. *PLAG1* has been identified as a cooperating neoplastic “hit” via a mutagenesis screen in *Cbfb-MYH11* murine leukemias and is upregulated in 20% of human AML^24^. Recent studies have reported a role for PLAG1 in stimulating expression of Mushashi-2, a protein that stimulates HSPC expansion^30^, and in expanding engrafting HSPC *ex vivo*(Hope et al, ASH 2017). Detection of abundant aberrant splicing and LV-PLAG1 mRNA fusions corroborates previous reports of LV-associated aberrant splicing ^8,33,34,51,52^. Of note, the LV-*PLAG1* splicing abrogated expression of non-translated E1 and E2, while maintaining expression of exons 3, 4 and 5, encompassing all translated sequences. We detected transcripts both with and without E4, predicting translation of both the short and long isoforms of PLAG1 protein^26^. Functional differences between the two PLAG1 isoforms are poorly understood.

We observed upregulation of expression of known downstream targets of PLAG1 in ZL34 cells, most notably *IGF2* and *MSI2*^30^. Imprinting-related downregulation of *IGF2* at the *IGF2/H19* locus maintains HSPC quiescence, with loss of imprinting resulting in murine HSPC proliferation, and exhaustion^53^. *MSI2* encodes the Musashi-2 RNA-binding protein, which promotes the expansion of long-term repopulating HSCs^31^ and regulates self-renewal^54^. Overexpression of *PLAG1* in CD34+ cells has been reported to stimulate HSPC expansion *ex vivo*(Hope et al, ASH 2017). It is interesting to note that *HMGA2* was downregulated in ZL34 hematopoietic cells compared to those from normal controls. *HMGA2 is* a transcription factor which functions as a critical upstream regulator of *PLAG1* expression^32^. An LV insertion resulting in overexpression of *HMGA2* was linked to the marked clonal expansion in an LV gene therapy trial noted above^8^ and based on the HSPC expansion seen in our animal, it seems possible that dysregulation of PLAG1 and downstream targets may have contributed to the expansion, although to our knowledge, no aberrant hematologic phenotype developed in that patient. The downregulation of *HMGA2* in ZL34 HSPC coincident with upregulation of PLAG1 and downstream targets suggests that HMGA2 may be involved in normal hematopoiesis, and regulated by feedback from downstream targets.

Although we suspect that genotoxicity linked to the *PLAG1* insertion likely played a key role in ZL34’s phenotype, we cannot rule out roles for other insertions. Given doses transplanted, HSPC with LV insertions in *PLAG1* have almost certainly been delivered to our other macaques without development of abnormal clonal hematopoiesis. An insertion just downstream of *KITLG* resulted in upregulation of SCF, with a small increase in serum SCF. Cell-associated SCF protein was increased in HSPC also expressing the KIT receptor, suggesting autocrine signaling, previously shown for other cytokines to drive proliferation in hematopoietic cells^55^. We have administered pharmacologic doses of SCF to macaques and have not observed eosinophilia or release of nRBC into the circulation, thus overexpression of SCF was unlikely to be the sole driver of ZL34’s aberrant hematopoiesis. However, it is of interest to note that production of fetal hemoglobin increases in baboons administered SCF^56^, and in human erythroid cultures supplemented with SCF^57^, suggesting dysregulation of SCF may have underlied elevated HbF in ZL34. Insertions upregulating expression or leading to aberrant splicing of *NCAM2*, *ISOC1*, or *EIF3E* seem less likely to be contributing to clonal expansion and the phentoype, given the lack of known roles for these genes in hematopoiesis or leukemogenesis. Expression of testin(*TES*) RNA was decreased, a gene with tumor suppressor activities, potentially via inhibition of cell migration and metastasis^58^. It is intriguing to speculate that loss of TES expression might have contributed to abnormal release of nRBC into the blood. The constraints of the primate model and our inability to derive immortalized lines from ZL34 precluded further direct investigations of the contribution of each specific insertion to the hematologic phenotype.

In conclusion, this fatal genotoxic clonal expansion adverse event linked to LV insertions is concerning. The implications of this event must be balanced against the encouraging safety and efficacy of LV HSPC gene therapy clinical trials in patients with life-threatening diseases. It is also important to note that the vector resulting in toxicity in our macaque model included a strong viral promoter/enhancer within the LTR, a design shown associated with increased genotoxicity in preclinical models, and not utilized in any clinical trial to date. However, we believe our findings imply that strong viral promoters should be avoided in LV whenever possible. In addition, approaches to decrease LV-related aberrant splicing should be pursued, given our evidence that upregulation of *PLAG1* was driven by primarily by hybrid transcripts. Targeted gene gene editing approaches avoid semi-random vector insertion and have the potential to be a less genotoxic technology, though these approaches are in their infancy and may be associated with a different set of off-target genotoxicities^59-61^.

## Acknowledgements

This research was supported by the NHLBI Division of Intramural Research. Di Yang was funded by the Scientific Research Training Program for Young Talents sponsored by Union Hospital, Tongji Medical College, Huazhong University of Science and Technology. We thank Keyvan Keyvanfar, the NHLBI FACS and DNA Sequencing and Genomics Cores, the NIH Biowulf High-Performance Computing Resource, NIH veterinary pathology, and NHLBI animal care staff for assistance. We acknowledge Patrick Duffy and Amber Raja for supplying marrow from animal CA3K. The authors have no relevant conflicts of interest to disclose.

## Author Contributions

Conceptualization: CED and CW; Analytics: DAE, SFC, LLT; Investigation: DAE, XF, DY, CW, KRC, IY, SD, XW, SSD; Resources: RL, HLM, NU, JFT, KH; Animal support: SGH, AK, MM, RED; Writing: CED, DAE, CW; Supervision: CED, CW, JFT.

## Supplemental Methods

### Cell purification and FACS analysis

PB and BM samples were separated into granulocyte and mononuclear cell fractions via centrifugation over lymphocyte separation medium (MP Biomedical) and stained with antibodies (Table S7) prior to flow cytometric analysis and sorting to purities >98%. CD34+ cells were purified by magnetic bead immunoselection (Miltenyi Biotec) to >95% purity. Neutrophils and eosinophils within the granulocyte pellet were separated based on CD33 expression. CD45^−^CD71^+^ flow gating was used to purify nRBCs for all figures with the exception of those requiring intracellular PLAG1/NCAM2/SCF FACS staining (Figures 4B; S9), in which solely CD45^**-**^ gating (>85% nRBC) was used.

### Colony forming assays

100 CD34^+^ BM cells purified from ZL34 BM were plated in 1ml methylcellulose media (Stemcell Technologies, Cat#: H4435) and cultured for 14 days. Single widely separated colonies were plucked for DNA extraction in 20uL DirectPCR Lysis Reagent buffer (Viagen Biotech, Cat# 401-E) with 1uL Proteinase K and 2mg/mL RNase.

### Vector integration site retrieval

Vector integration sites were identified retrieved as described^50^. Briefly, genomic DNA was sheared (Covaris) to an average size of 300-500bp, end repaired and dA-tailed. T-Linkers (Oligoseq) were ligated to the resulting fragments. PCR was performed using one LTR-specific primer and one linker-specific primer followed by nested primers containing Illumina sequencing adaptors. The resulting library was sequenced on an Illumina MiSeq. Integration site junctions were trimmed with custom Perl scripts and mapped to rheMac8 using Blat.

### Analysis of vector integration sites relative to known regulatory sequences

The locations of vector integration sites were compared to CTCF binding sites as determined by chromatin immunoprecipitation sequencing (ChIP-seq). FASTQ files from four ChIP-seq experiments on the hepatocytes of three male macaques were obtained from ArrayExpress (accession number E-MTAB-437)^62^. We used STAR^63^ to map the reads to the rheMac8/Mmu8_8.0.1 reference genome and MACS2^64^ to predict the genomic locations of CTCF binding sites from the aligned reads. Finally the Bioconductor package GenomicRanges^65^ was used to compute overlaps between vector integration sites and CTCF binding sites.

Locations of vector integration sites were also more broadly compared to the locations of other regulatory elements as determined by an assay for transposase-accessible chromatin using sequencing (ATAC-seq) data. ATAC-seq data from 18 human hematopoietic lineages were downloaded from the Gene Expression Omnibus (GEO accession number GSE96772)^66^. We translated the ATAC-seq peaks from human genomic coordinates (aligned to hg19) into macaque genomic coordinates (aligned to rehMac8) using the UCSC batch coordinate conversion program, liftover, using the appropriate chain file from the USCS Genome Browser website (http://genome.ucsc.edu). Again we used the Bioconductor package GenomicRanges^65^ to compute overlaps between vector integration sites and ATAC-seq peaks.

### Hemoglobin chain measurements

Whole blood cell pellets were lysed in 100 µL HPLC-grade water by pulse-vortexing for 30 seconds (3X) followed by one cycle of freeze-thawing and centrifugation at 16,000Xg for 15 minutes. 10 µL of 100mM TCEP (Thermo Fisher Scientific) was added, and after 5 minutes incubation at room temperature, 85µl of 0.1% TFA/32% acetonitrile was added. 10μL samples were analyzed at a 0.7 ml/minute flow rate for 50 minutes using the Agilent 1100 HPLC (Agilent Technologies) equipped with the Aeris 3.6μm Widepore C4 200 (250×4.6mm, Phenomenex, Torrance, CA) reverse phase column, using solvent A; 0.12% TFA in water and solvent B, 0.08% TFA in acetonitrile. A starting concentration of solvent B was designated as 35% for the separation of globin proteins and % of solvent B was changed as follows: 3 minutes at up to 41.2%, 3 minutes at up to 41.6%, 5 minutes at up to 42%, 4 minutes at up to 42.4%, 6 minutes at up to 42.8%, 6 minutes at up to 44.4%, 6 minutes at up to 47%, 7 minutes at up to 75% and re-equilibrated for 10 minutes at 35%. The globin types were detected at 215nm and confirmed by an Agilent HPLC-6224 mass spectrometer equipped with an ESI interface and a time-of-flight (TOF) mass detector as previously described ^67,68^.

### Soluble SCF detection by using ELISA

Plasma was obtained by centrifuging whole blood at 2000g for 15 minutes at 4°C. 1 * 10^7^ PB MNC were cultured for 24 and 48 hours in RPMI10 media supplemented with 10% fetal bovine serum. The stem cell factor human SimpleStep ELISA Kit (Abcam, Cambridge, MA, Cat#176109) was used to measure SCF levels in plasma and conditioned culture media.

### RNA sequencing and analysis

CD45^−^CD71+ nRBC and CD34^+^ cells from ZL34(samples collected days 410-540 post-transplant) or control monkey PB or BM(4 age-matched untransplanted animals, and 1 transplanted animal day 465 post-transplant) were purified by FACS sorting (nRBC) or immunoselection (CD34^+^ cells). Total RNA was extracted with RNAzol RT(MRC, OH, USA). RNA libraries were prepared using the Illumina TruSeq Stranded Total RNA kit with Ribo-Zero and sequenced on an Illumina HiSeq 3000.

RNA-seq data analysis was performed using STAR ^63^, DESeq2 ^69^, and custom R code. Genome assembly Mmul_8.0.1 was used. Macaque annotation from Ensembl release 92 was used. Analysis of nRBC and HSPC data together was performed in DESeq2 using the design formula ~ source + celltype + condition, where source was PB or BM, celltype was HPSC or nRBC, and condition was ZL34 or control. nRBC-only analysis was performed using the design formula ~ source + condition. Built-in Cook’s cutoff was used in DESeq2 to alleviate the effect of outliers on resulting differentially expressed genes.

### Histologic and immunohistochemical analyses

BM biopsies were obtained from posterior iliac crests. Following fixation and decalcification, sections were prepared. Spleen samples were obtained at time of autopsy. Antibodies utilized for immunohistochemical staining are listed in Table S7.

### Analysis of differential usage of splice variants

We implemented a pipeline to detect fusion genes in parallel with computing differential usage of gene features and detects. In brief, STAR v 2.6.0c ^63^ was used to align reads to a combination *Macaca mulatta*, (ENSEMBL Mmul_8.0.1) plus custom vector reference genomes and a combination *Macaca mulatta* annotation (ENSEMBL, Mmul_8.0.1.92 general transfer format) plus custom annotation for the vector sequence. Then STAR-Fusion (https://github.com/STAR-Fusion/STAR-Fusion/wiki) was used to compute fusion genes using a custom genome resource library for the macaque and vector reference genomes, which was built using FusionFilter. Differential usage of gene features was computed using QoRTs^70^ and JunctionSeq^71^. Our implementation of differential gene expression relied on DESeq2^69^. All components of our pipeline were unified under custom shell and R scripts that were run on the NIH High Performance Computing cluster. All of our source code and scripts is available via BitBucket (https://bitbucket.org/DunbarLab_Releases/variant-splicing_fusion-genes).

### Data availability

All data and code used in this study (RNA-seq and barcode data) will be made available upon request from the corresponding authors.

---

**Figure S1.**
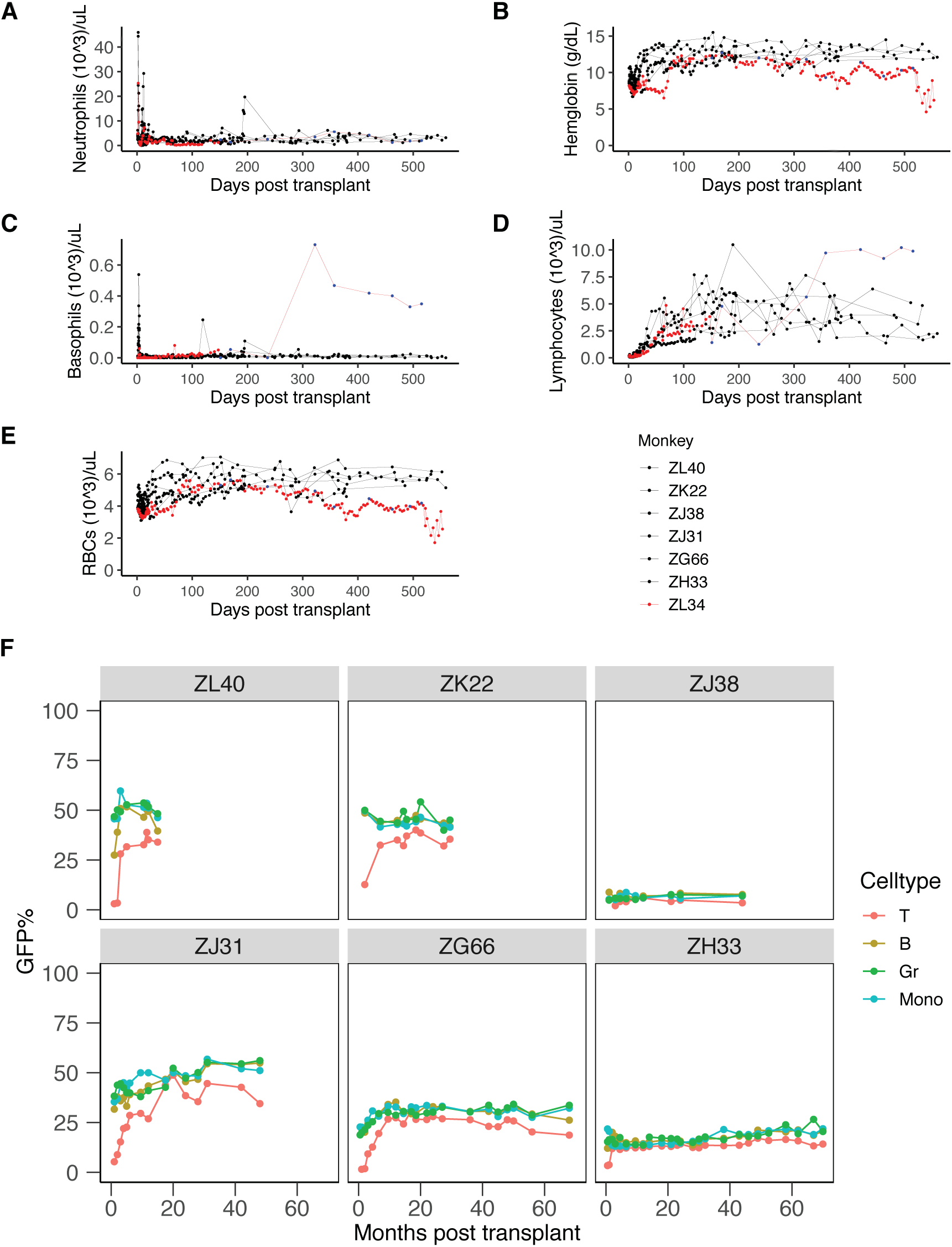
CBC counts in ZL34 and controls post-transplant. Neutrophil counts **(A)**, hemoglobin levels **(B),** basophil counts **(C)**, lymphocyte counts **(D)**, and red blood cell counts **(E)** post-transplant as determined by standard CBC (red or black dots) or automated visual scanning (blue dots) for ZL34 and 6 control macaques transplanted with HSPC transduced with the same MSCV LV vector. **(F)** GFP-positivity of T, B, Monocyte (Mono), and Granulocyte (Gr) subsets in 6 control macaques transplanted with HSPC transduced with the same MSCV vector as ZL34.

**Figure S2.**
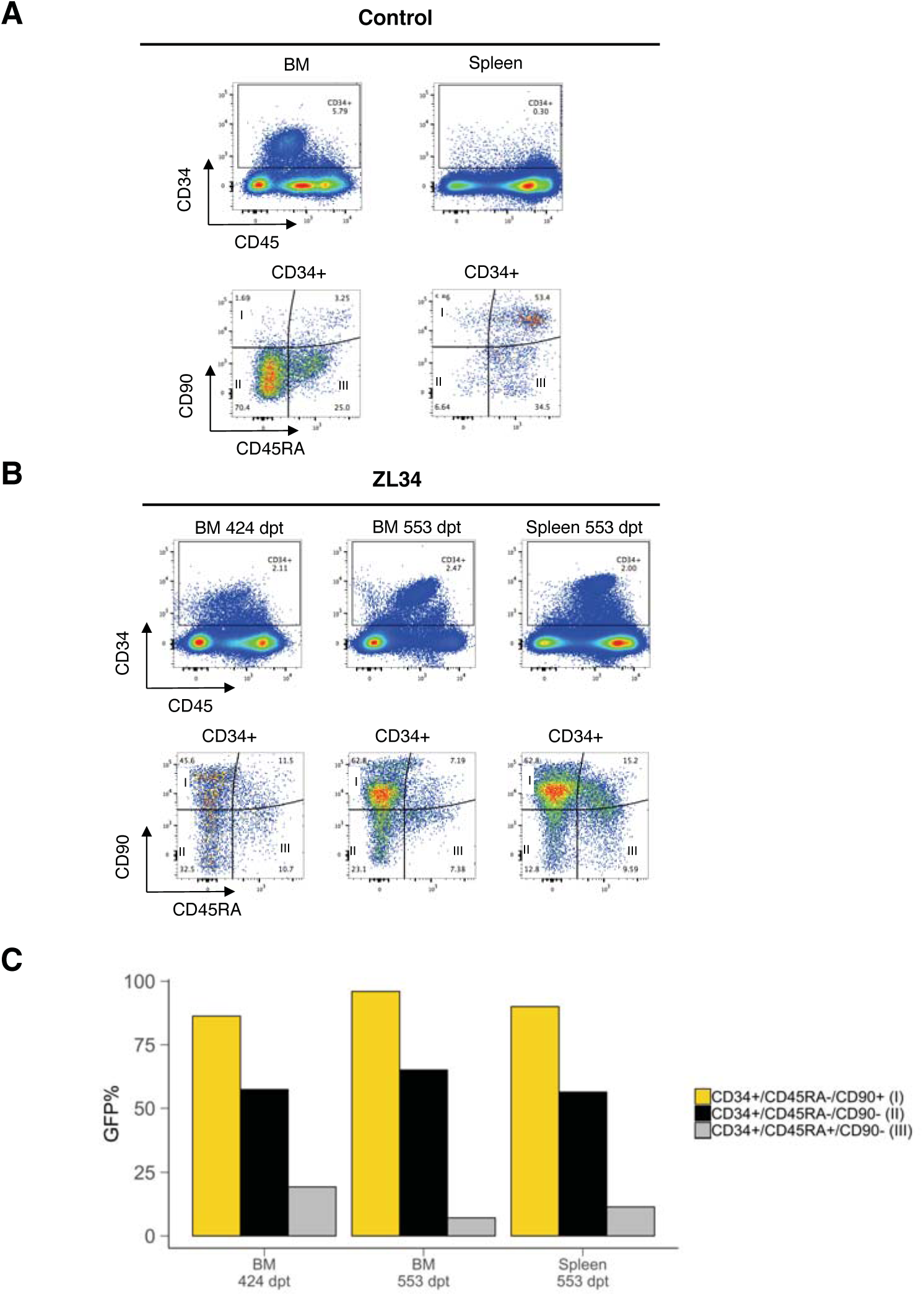
Counts and GFP% of BM and 5pleen CD34 subsets in ZL34 and controls. CD34+ cells were extracted from BM and Spleen in a control macaque **(A)** and for two time points for ZL34 **(B)**, and evaluated for expression of CD34, CD45RA, and CD90. CD34+/CD45RA-/CD90+ (I) cells represent an HSC-enriched fraction, while CD34+/CD45RA-/CD90-(II) and CD34+/CD45RA+/CD90-(III) represent more mature progenitors, as defined^20^. **(C)** GFP-positivity determined by flow cytometry for subsets I, II, and III.

**Figure S3.**
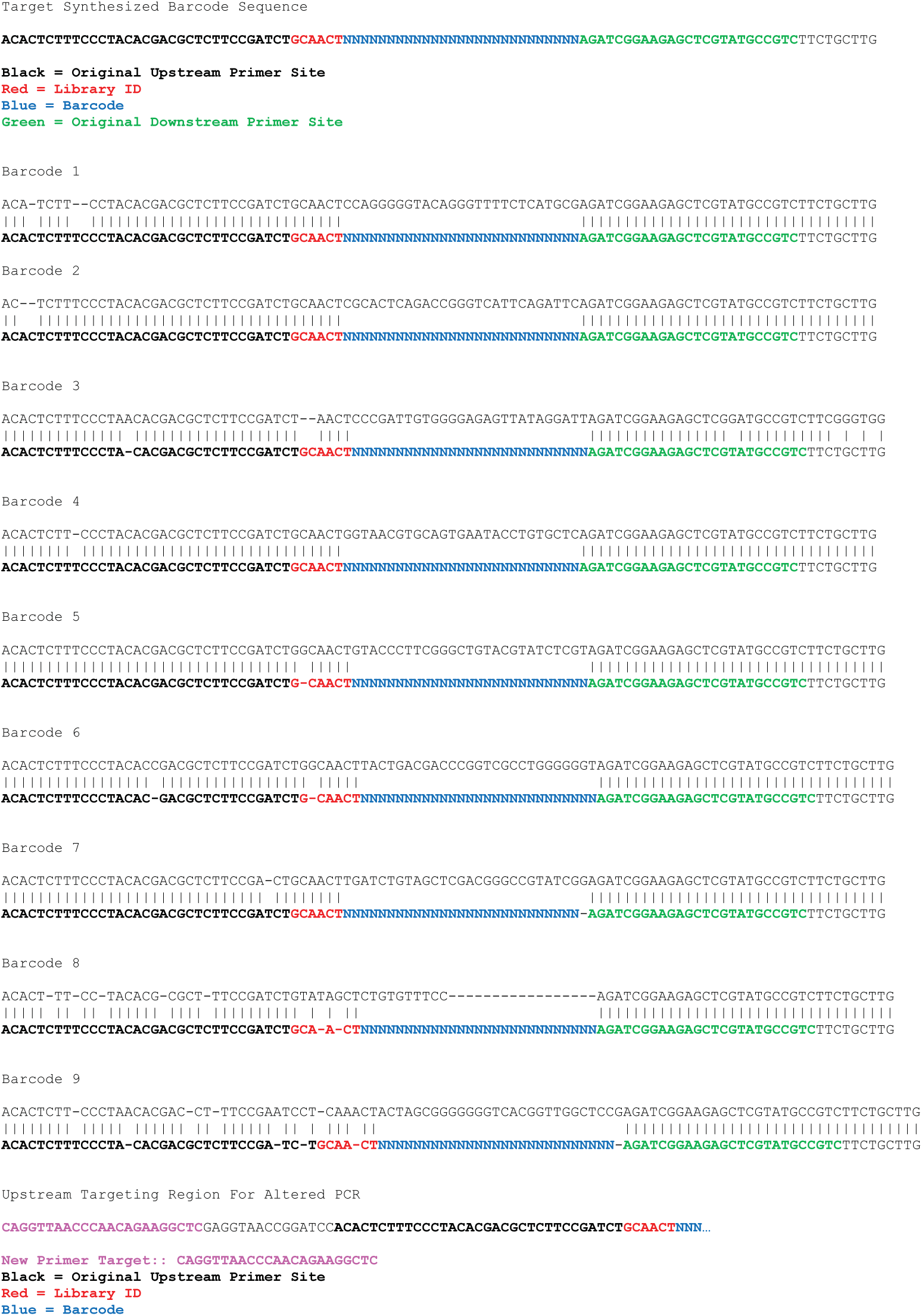
Sequence and mutations of 9 barcodes recovered from ZL34 nRBC DNA. Laboratory standard barcode recovery protocol and an optimized barcode recovery protocol recovered 9 barcodes within ZL34’s nRBC DNA, 3 of which (#7-9), contained mutations in the forward primer/read 1 primer site prohibitive to efficient PCR and/or Illumina sequencing using the standard primers. Upstream targeting region for the optimized altered PCR is shown.

**Figure S4:**
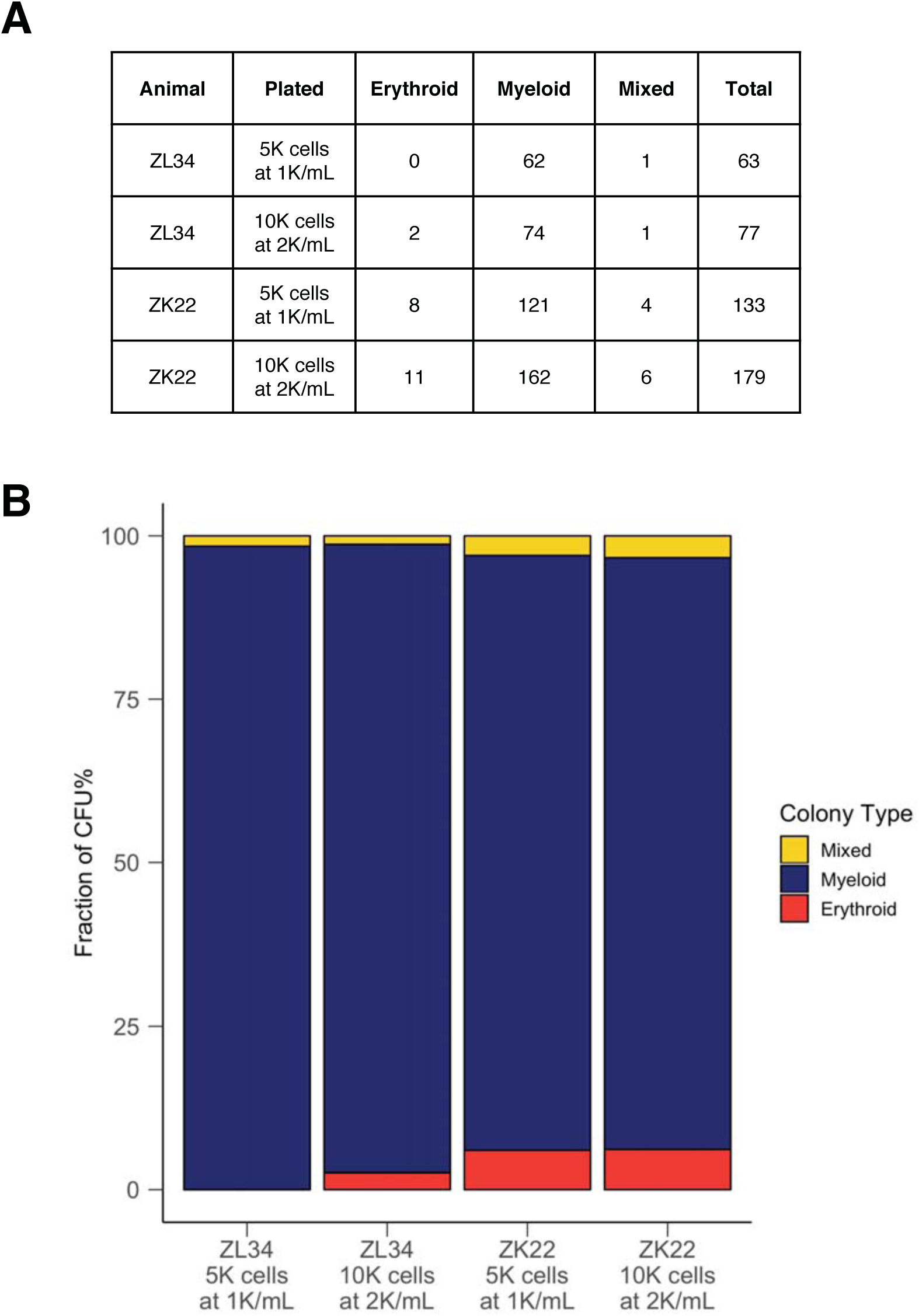
ZL34 versus control CFU analysis. Bone marrow CD34+ cells purified from ZL34 at day 305 post-transplantation and a control macaque ZK22 day 354 post-transplantation were plated for CFU enumeration. **(A)** Counting of erythroid, myeloid and mixed CFU 14 days after plating in ZL34 versus ZK22 for 1000 or 2000 cells plated/mL. **(B)** Proportion of myeloid, erythroid, and mixed CFU in each sample.

**Figure S5.**
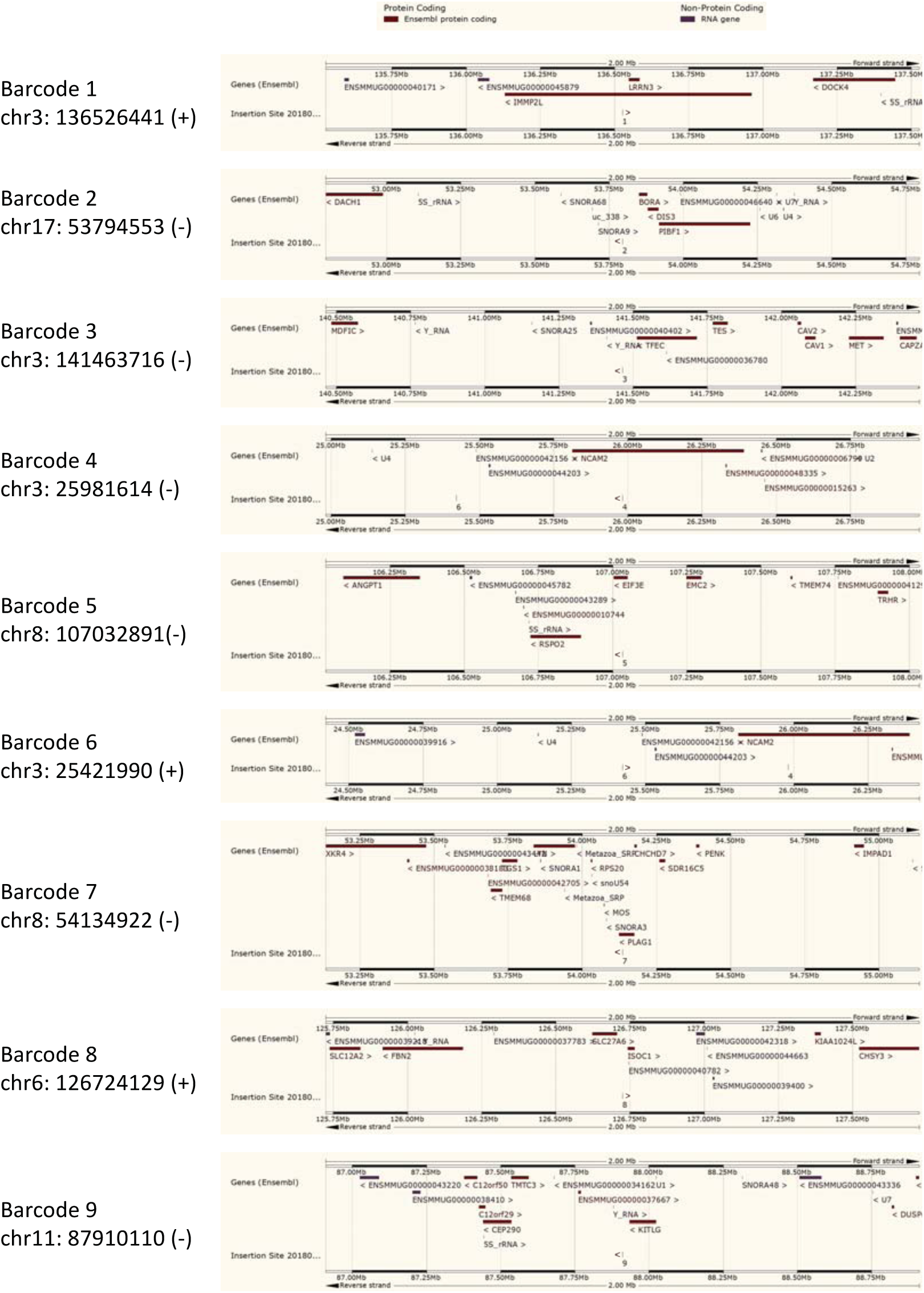
Genomic location of the 9 insertions. Genomic coordinates of insertion sites for the 9 lentiviral insertions are plotted in 2 Mb windows on the rhesus macaque genome (Mmul8) in the Ensembl genome browser showing the proximal features and genes associated with the insertions.

**Figure S6.**
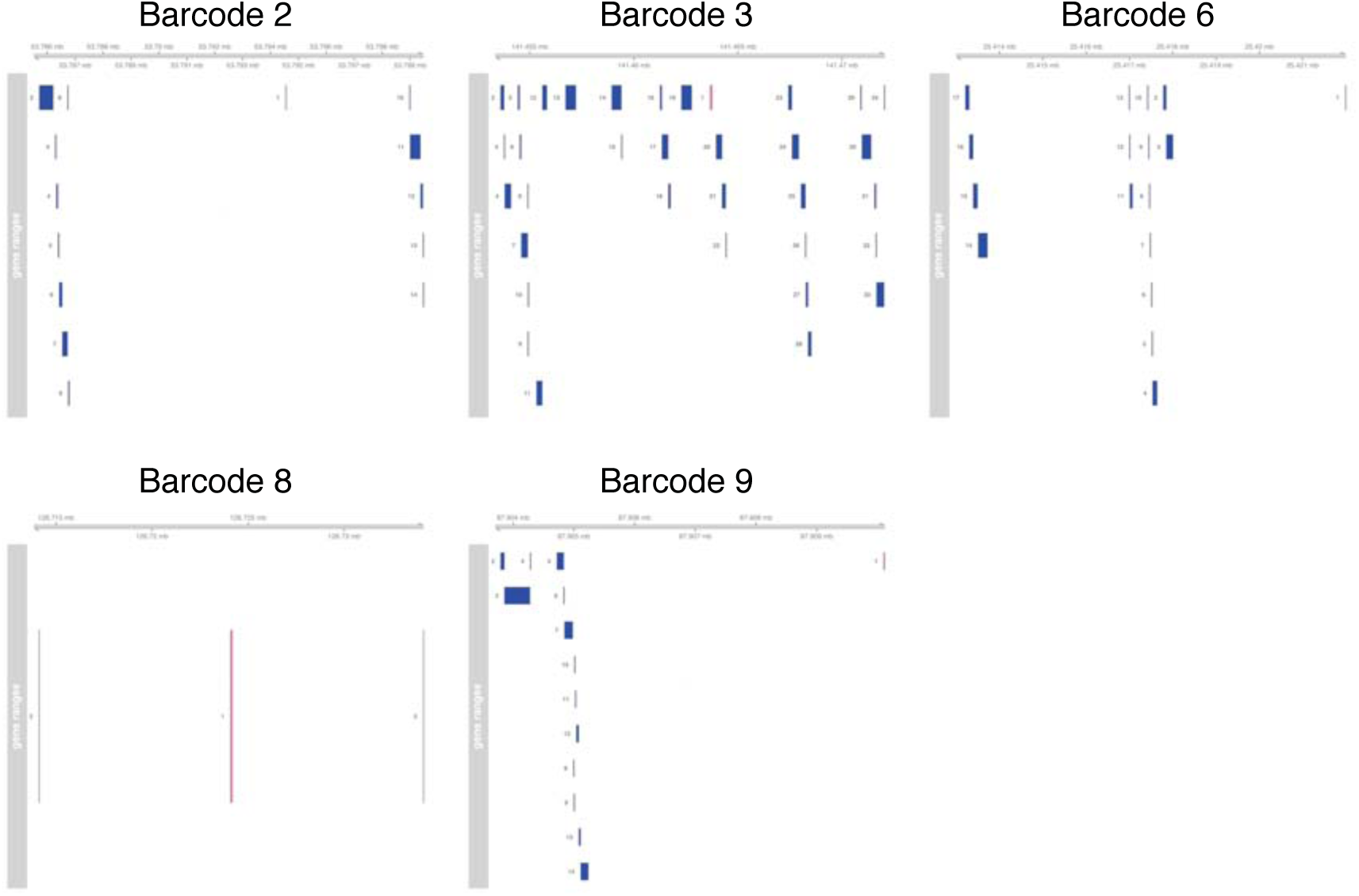
Regulatory element proximity to Intergenic insertions. Depiction of the genomic neighborhood of intergenic vector insertions relative to putative regulatory elements as determined by ATAC-seq data from 18 human hematopoietic lineages. Genomic regions depicted in hot pink represent the vector insertion; genomic regions depicted in blue represent ATAC-seq peaks that were converted from human to macaque genomic coordinates as described in the supplemental methods.

**Figure S7.**
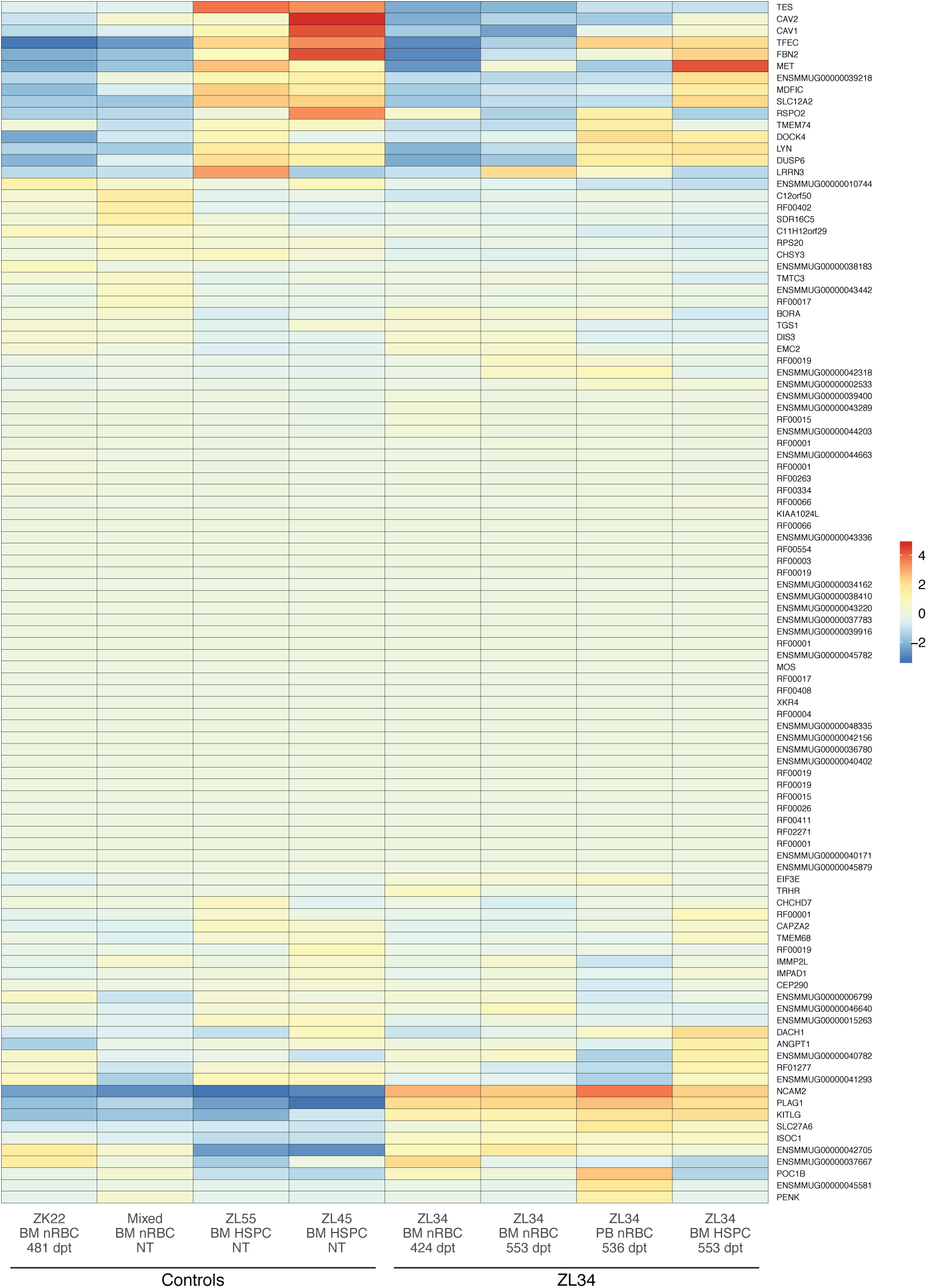
Mean-centered gene expression levels of all genes and features proximal to the 9 insertions. Figure 3 illustrates the significant dysregulation of only 5 of these genes (*PLAG1, KITLG, NCAM2, ISOC1,* and *TES*) when comparing nRBC plus HSPC samples in ZL34 and controls. Expression levels are rlog values from DESeq2.

**Figure S8.**
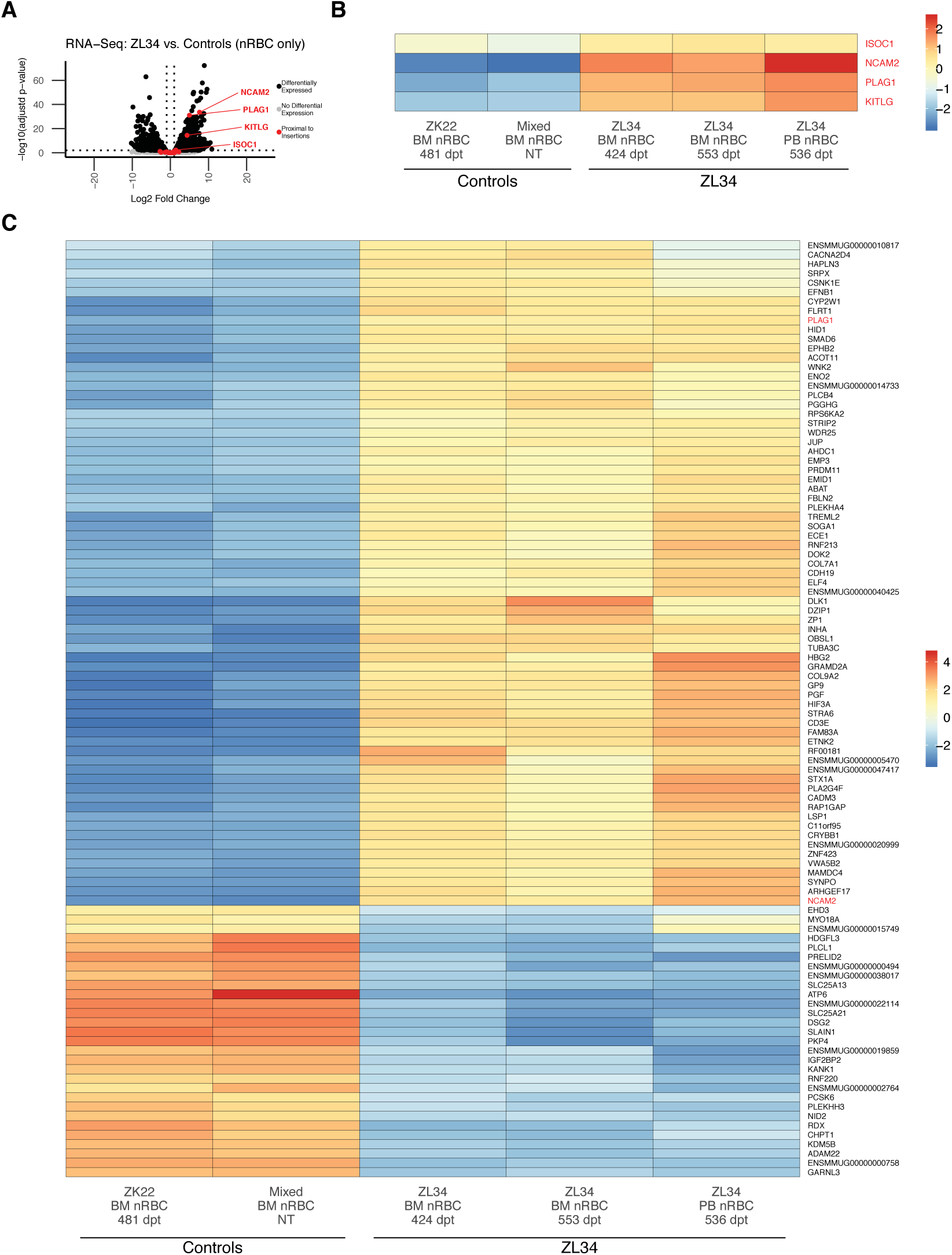
RNA-seq analysis restricted to nRBC samples. **(A)** Volcano plot of RNA-seq data restricting analysis to solely nRBC samples from Figure 3, showing significant upregulation of only 4 insertion-proximal (within 1 Mb) genes: *PLAG1, NCAM2, KITLG, and ISOC1.* **(B)** Mean-centered expression levels of *NCAM2, KITLG, PLAG1, and ISOC1*, 4 differentially expressed genes each within 1 Mb of a different insertion, shown across all nRBC RNA-seq samples. Expression levels are rlog values from DESeq2. **(C)** Mean-centered expression levels of the top 100 differentially expressed genes in ZL34 compared to controls, when analysis is restricted to nRBC samples only. Expression levels are rlog values from DESeq2.

**Figure S9.**
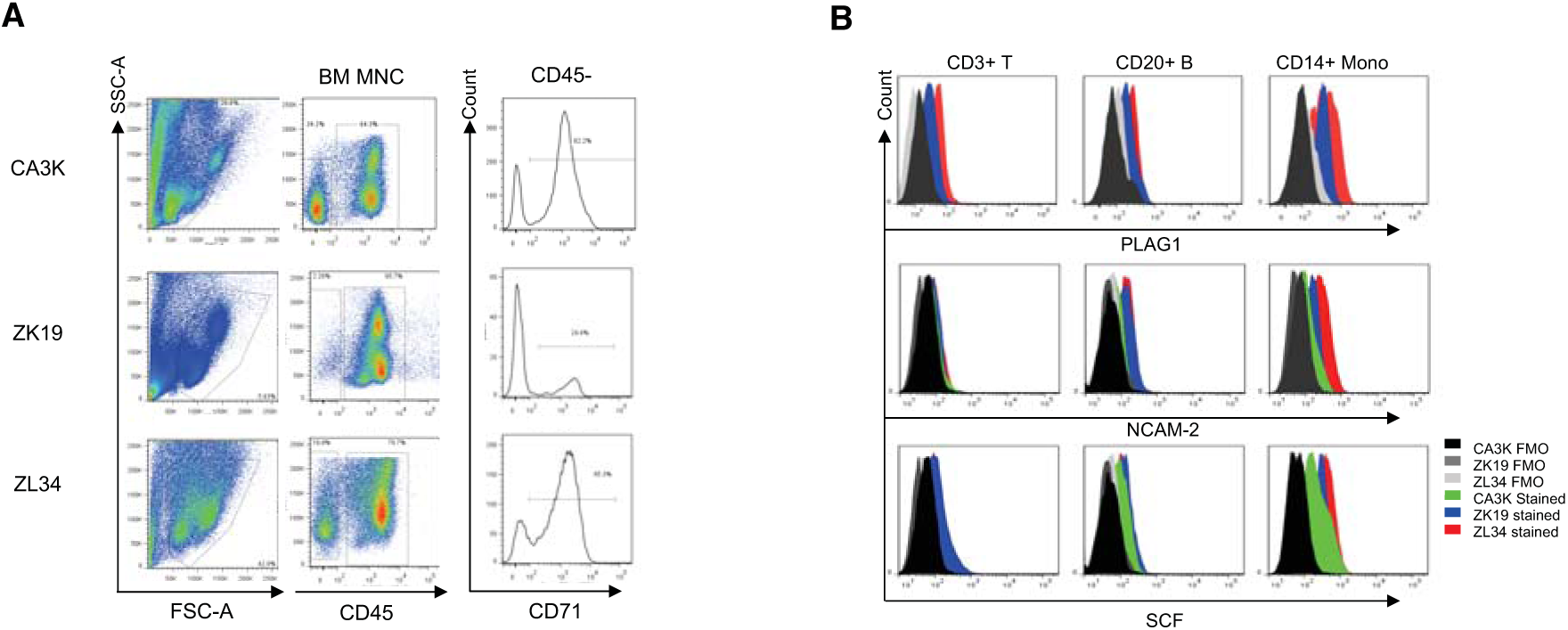
Aberrant protein expression of PLAG1, NCAM2 and 5CF in ZL34 hematopoietic cells. **(A)** CD71 expression on CD45-cells in a control monkey (ZK19), a monkey recovering from malarial anemia (CA3K) and ZL34, documenting expanded erythroid CD45-cells in ZL34. **(B)** Flow cytometry analysis of the PLAG1, NCAM2 and SCF expression on T cells, B cells and monocytes in ZK19, CA3K, and ZL34.

**Figure S10.**
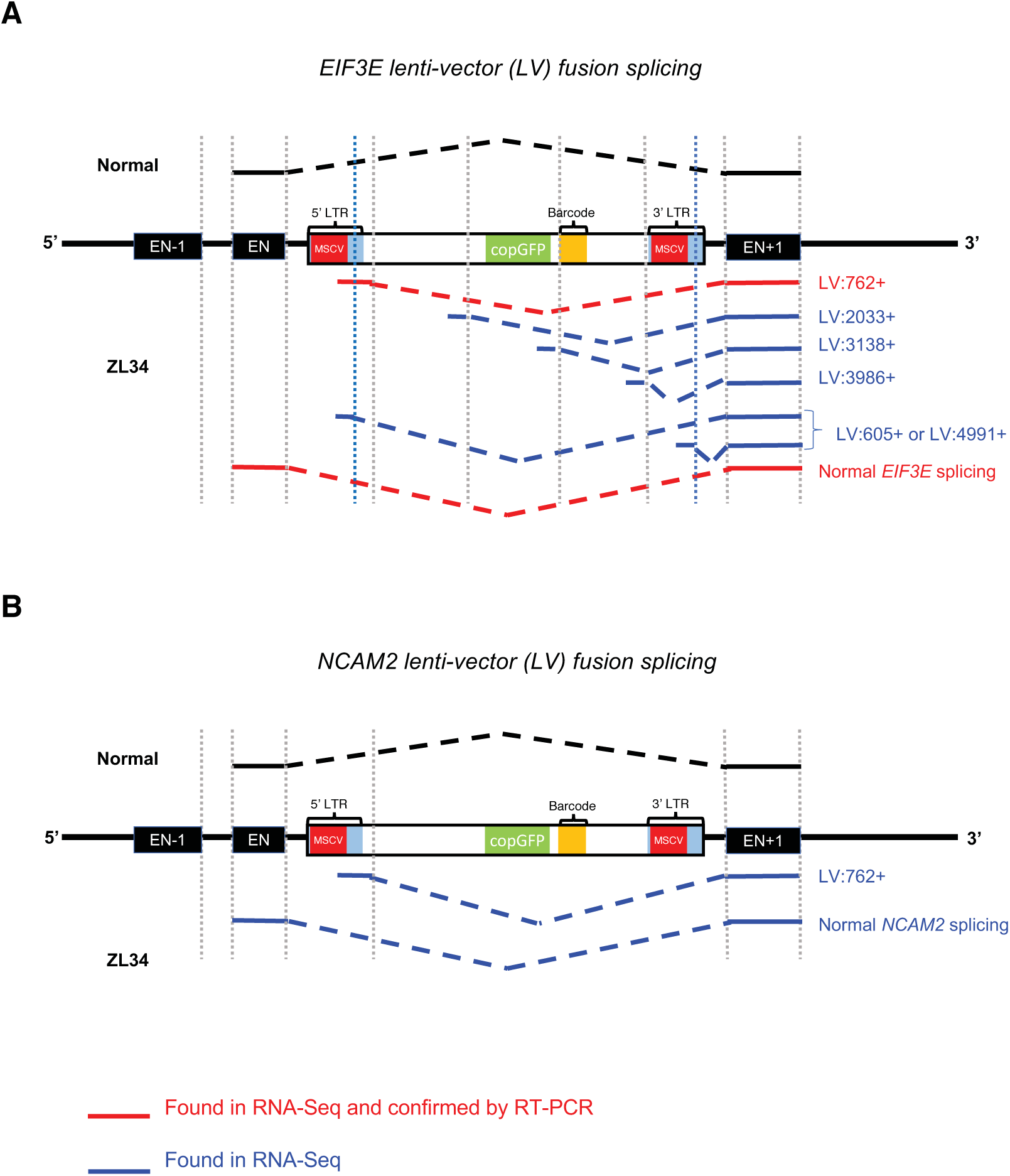
*EIF3E* and *NCAM2* hybrid transcript detection. Hybrid vector-gene transcripts were detected within ZL34 for Barcode 5 and its intronic host, *EIF3E* **(A)**, as well as for Barcode 4 and its intronic host, *NCAM2* **(B)**. Subsets of the results were able to be confirmed by RT-PCR and Sanger sequencing.

## Supplemental Tables

### See accompanying Excel files for

**Table S1. Transplant information and follow-up for all transplanted letivirally-barcoded macaques.**

**Table S2. Viral integration site retrieval in ZL34.** Viral integration site retrieval (5’ and 3’-based) in ZL34 nRBC DNA at 309 and 393 days post-transplant, and in granulocyte DNA at 96 days post-transplant. Green denotes the integration site recovered has been matched to one of the originally recovered barcodes and is concluded to arise from the single expanded clone in ZL34.

**Table S3. All RNA-seq differentially expressed genes for H5PC+nRBC comparison and nRBC-only comparisons.** Differentially expressed genes reported from DESeq2, identified as described in Bulk RNA-seq analysis. ens_id = Ensembl gene ID. baseMean = the average of the DESeq2 normalized count values for all samples, normalized for sequencing depth. log2FoldChange = DESeq2 estimated effect size in log2 scale, comparing ZL34 to controls. lfcSE = standard error of the log2FoldChange estimate. stat = DESeq2 Wald statistic. pvalue = Wald test p-value. padj = Benjamini-Hochberg adjusted p-value.

**Table S4. Differentially spliced genes in ZL34.** Differential expression of gene features (e.g. exons or exon junctions) obtained from the comparison of RNA-seq data from (i) four samples from ZL-34 (1 sample of nucleated erythrocytes from peripheral blood, 2 samples of nucleated erythrocytes from bone marrow and 1 sample of CD34+ cells obtained from bone marrow) and (ii) 4 samples from a wild type macaque (2 samples of nucleated erythrocytes obtained from bone marrow and 2 samples of CD34+ HSPCs obtained from BM). Differential expression of features was computed with our pipeline and a custom index for the combined macaque and lentiviral as described in the supplemental methods. Tab 1 is a gene level overview of features (e.g. exons or junctions) that are differentially expressed with an adjusted p-value of less than 0.05. The meanings of the columns is described in comments added to each column and also tabulated below. Tab 2 is a more detailed presentation of the results at the level of individual gene features. Again, the meanings of each of the columns is described in comments added to each column and also tabulated below.

Columns on Tab 1:

Column 1 (ID):

ENSEMBL gene ID.(Macaque ENSEMBL release 92)

Column 2 (Gene Symbol):

HGNC symbol corresponding to ENSEMBL ID, if known

Column 3 (Description):

Description of gene function, if known.

Column 4 (Chr):

Chromosome on which gene is located.

Column 5 (Start):

(1-based) position of the start of gene

Column 6 (End):

(1-based) end of the gene.

Column 7 (Strand):

Strand on which gene is located.

Column 8 (baseMean):

The base mean normalized coverage counts for the locus across all conditions.

Column 9 (geneWisePadj):

The gene-level p-value that one or more features belonging to this gene are differentially used. This value will be the same for all features belonging to the same gene.

Column 10 (mostSIgID):

The sub-feature OD for the most significant exon or splice junction belonging to the gene.

Column 11 (mostSIgPadj):

The adjusted p-value for the most signifiance exon or splice-junction belonging to the gene.

Column 12 (numExons):

The number of known non-overlapping exonic regions belonging to the gene.

Column 13 (numKnown):

The number of known splice junctions belonging to the gene.

Column 14 (numNovel):

The number of novel splice junctions belonging to the gene.

Column 15 (exonsSig):

The number of statistically significant non-overlapping exonic regions belonging to the gene.

Column 16 (knownSIg):

The number of statistically significant known splice junctions belonging to the gene

Column 17 (novelSig):

The number of statistically significant novel splice junctions belonging to the gene.

Column 18 (numFeatures):

The columns numExons, numKnown, and numNovel, separated by slashes.

Column 19 (numSig):

The columns exonsSig, knownSIg, and novelSig, separated by slashes.

Columns on Tab 2:

Column 1 (ID):

ENSEMBL gene ID.(Macaque ENSEMBL release 92)

Column 2 (testable):

Whether enough reads to enable statistical comparison.

Column 3 (pvalue):

P-value for differential expression of the gene of which this is feature

Column 4 (padjust):

Adjusted p-value of the gene of which this is feature.

Column 5 (Chr):

Chromosome on which gene is located.

Column 6 (Start):

(1-based) position of the start of gene.

Column 7 (End):

(1-based) end of the gene.

Column 8 (Strand):

Strand on which gene is located.

Column 9 (transcripts):

Known transcripts involving this feature.

Column 10 (featureType):

Type of feature.

Column 11 (p-adj):

Adjusted p-value for the test of differential usage.

Column 12 (log2FC(ZL34/WT)):

Log 2 fold change for ZL34 versus WT.

**Table S5. Fusion LV-endogenous gene detection in ZL34.** Table of lentiviral-endogenous mRNA fusions found in RNA-seq data obtained from four samples from ZL-34 (1 sample of nucleated erythrocytes from peripheral blood, 2 samples of nucleated erythrocytes from bone marrow and 1 sample of CD34+ cells obtained from bone marrow using our pipeline as described in the supplemental methods). The first column tabulates the left break point which is located in the lentiviral insertion (LVI). The LVI act as a splice donor and five different (LVI) break points were observed in fusion genes with EIF3E; 3 of the same LVI breakpoints were also observed in fusions with PLAG1 and NCAM2; we highlight the fact that the other 2 breakpoints were not observed with rows with zero entries. The second column depicts the right break point. In each case these are starts of known exons. Columns 3 through 6 denote the fusion fragments per million as computed from our pipeline and described in the supplemental methods. Most of the fusion genes were confirmed by PCR as indicated in the 7th column of the table. As depicted in the 8th column two of the splice junctions were previously reported by Cesana et al^33^.

**Table S6. Primers used for barcode retrieval.**

**Table S7. Antibody information**.

